# Non-uniform distribution of dendritic nonlinearities differentially engages thalamostriatal and corticostriatal inputs onto cholinergic interneurons

**DOI:** 10.1101/2021.11.29.470423

**Authors:** Osnat Oz, Lior Matityahu, Aviv Mizrahi-Kliger, Alexander Kaplan, Noa Berkowitz, Lior Tiroshi, Hagai Bergman, Joshua A. Goldberg

## Abstract

The tonic activity of striatal cholinergic interneurons (CINs) is modified differentially by their afferent inputs. Although their unitary synaptic currents are identical, in most CINs cortical inputs onto distal dendrites only weakly entrain them, whereas proximal thalamic inputs trigger abrupt pauses in discharge in response to salient external stimuli. To test whether the dendritic expression of the active conductances that drive autonomous discharge contribute to the CINs’ capacity to dissociate cortical from thalamic inputs, we used an optogenetics-based method to quantify dendritic excitability. We found that the persistent sodium (NaP) current gave rise to dendritic boosting, and that the hyperpolarization-activated cyclic nucleotide-gated (HCN) current gave rise to a subhertz membrane resonance. This resonance may underlie our novel finding of an association between CIN pauses and internally-generated slow wave events in sleeping non-human primates. Moreover, our method indicated that dendritic NaP and HCN currents were preferentially expressed in proximal dendrites. We validated the non-uniform distribution of NaP currents pharmacologically; with two-photon imaging of dendritic back-propagating action potentials; and by demonstrating boosting of thalamic, but not cortical, inputs by NaP currents. Thus, the localization of active dendritic conductances in CIN dendrites mirrors the spatial distribution of afferent terminals and may promote their differential responses to thalamic *vs*. cortical inputs.

## Introduction

The striatal cholinergic interneuron (CIN) is a key modulator of the striatal microcircuitry, impacting the neuronal excitability, synaptic transmission and synaptic plasticity of spiny projection neurons (SPNs), as well as other striatal interneurons (Abudukeyoumu et al., 2019; Assous, 2021; Goldberg et al., 2012; Matityahu et al., 2022). There are two processes that drive the ongoing release of acetylcholine (ACh) by CINs. First, the two glutamatergic inputs arising from cerebral cortex and intralaminar nuclei of the thalamus can drive CINs to discharge (Bradfield et al., 2013; Ding et al., 2010; Doig et al., 2014; Kosillo et al., 2016; Lapper and Bolam, 1992; Mamaligas et al., 2019; Matsumoto et al., 2001; Sharott et al., 2012; Thomas et al., 2000). While the unitary synaptic currents generated by these two inputs in CINs are identical (Aceves Buendia et al., 2019), thalamic inputs to CINs dominate in the sense that they that give rise to larger excitatory post-synaptic potentials (EPSPs) in acute striatal slices (Johansson and Silberberg, 2020) and can trigger abrupt pause responses, often flanked by excitatory peaks, to external saliency-related cues (Apicella et al., 1991; Goldberg and Reynolds, 2011; Graybiel et al., 1994; Kimura et al., 1984; Matsumoto et al., 2001; Morris et al., 2004; Raz et al., 1996). In contrast, cortical inputs to CINs are weaker in that, in acute striatal slices, they give rise to smaller EPSPs in most CINs (Mamaligas et al., 2019) and cannot trigger the pause response (Ding et al., 2010), although, these differences are less pronounced in intact animals (Doig et al., 2014).

Second, even in the absence of afferent input, CINs exhibit multiple autonomously generated discharge patterns – including regular and irregular pacemaking, as well as burst firing (Bennett et al., 2000; Bennett and Wilson, 1999; Goldberg and Reynolds, 2011; Goldberg and Wilson, 2010). These firing patterns are generated by an interplay between various nonlinear ionic currents including voltage- and calcium-activated K^+^ currents, as well as two voltage-dependent pacemaker currents: the hyperpolarization-activated cyclic nucleotide-gated (HCN) current; and the persistent Na^+^ (NaP) current (Bennett et al., 2000; Deng et al., 2007; Goldberg et al., 2009; Goldberg and Wilson, 2010, 2005; McGuirt et al., 2021; Oswald et al., 2009; Song and Surmeier, 1996; Wilson, 2005; Wilson and Goldberg, 2006). The main purpose of these pacemaker currents presumably is to guarantee the ongoing release of ACh onto the striatal microcircuitry by sustaining the autonomous discharge of CINs. Nevertheless, these pacemaker (and other subthreshold) currents will also impact how cortical and thalamic inputs are integrated by the CINs.

Attaining a mechanistic (e.g., dynamical systems) understanding of how the repertoire of CIN firing patterns is generated, requires a full characterization of the nonlinear properties of the pacemaker (and other) currents (e.g., by determining their voltage dependence and kinetics) – which is a daunting endeavor. In contrast, understanding how these currents impact synaptic inputs is a simpler task. Because individual synaptic inputs are small, the membrane nonlinearities can be *linearized* making the analysis of their impact simpler and more general – a treatment called the quasi-linear membrane approximation. This analysis dates back to Mauro (Koch, 1984; Mauro et al., 1970) and has shown that quasi-linear membranes can give rise to two qualitatively different transformations of inputs: *amplification* and *resonance* (Goldberg et al., 2007; Hutcheon and Yarom, 2000). Being a linear approximation, the quasi-linear approximation is amenable to Fourier analysis, which helps to better define these transformations as linear (time-invariant) filters on the input in frequency space.

Amplification arises from regenerative ionic currents – that provide positive feedback – including inward (depolarizing) currents activated by depolarization, such as the NaP current. Here the main effect in frequency space is amplification of the amplitude response (as compared to the response of a passive linear membrane). Resonance arises from restorative currents – that provide negative feedback – including inward currents activated by hyperpolarization, such the HCN current. In frequency space, the defining properties of resonance is a peak (at a non-zero frequency) in the amplitude response, and a zero-crossing of the phase delay (at a nearby frequency). On a practical level, the quasi-linear response properties of a membrane can be measured by providing a small-amplitude, sinusoidally-modulated voltage command to an intracellularly-recorded neuron and recording the resultant sinusoidal output current. Calculating the ratio of the voltage amplitude to the current response yields an estimate of the membrane’s impedance (which is, loosely speaking, an indication of the neuron’s “input resistance” to a sinusoidal input as a function of its frequency). Membrane impedances have been reported for various neuronal types in the brain (Hutcheon et al., 1996; Ulrich, 2002), including in the striatum (Beatty et al., 2015). Beatty and collaborators (2015) found that CINs exhibit a resonance in the vicinity of 1 Hz.

Moreover, they found that the shape of the impedance function depended on the holding voltage, which is understandable given that the amplitude of the various subthreshold currents is voltage-dependent. Finally, with use of tetrodotoxin (TTX), a selective antagonist of voltage-activated Na^+^ (Nav) channels, Beatty and collaborators (2015) demonstrated that NaP currents contributed to these filtering properties of the CIN membrane.

Use of somatic voltage perturbations, however, fails to discriminate the role played by the CINs’ dendritic arbor *per se* in transforming synaptic inputs. The CINs’ dendritic arbor can span well over half a millimeter from the soma (Wilson, 2004), with cortical inputs terminating on distal dendrites and thalamic inputs terminating perisomatically and on proximal dendrites (Doig et al., 2014; Lapper and Bolam, 1992; Mamaligas et al., 2019; Thomas et al., 2000). Both dendritic morphology (as taught by cable theory) and dendritic nonlinearities will lead to distal cortical inputs being integrated differently from proximal thalamic input. Thus, the impact of membrane nonlinearities on the quasi-linear approximation will depend on *where* they are expressed throughout the dendritic arbor. To address this question, we recently developed an optogenetics-based experimental method (that relies on the use of quasi-linear cable theory and Fourier analysis) to determine the duration of the delays introduced by dendrites, and how these delays impact the rapidity and fidelity of a neuron’s response to its input. We used our method to study GABAergic neurons of the substantia nigra pars reticulata (SNr) that expressed channelrhodopsin-2 (ChR2). We illuminated (with a 470 nm LED) either a small perisomatic region or the entire dendritic arbor. Comparison of the two illumination regimes enabled us to demonstrate that dendrites (that in the SNr can be >700 µm long) introduce a significant integration delay. The analysis also yielded that SNr dendrites behaved like passive linear filters, without evidence for amplification or resonances, or any dependence on holding voltage (Tiroshi and Goldberg, 2019).

In the present study, we apply our method to study nonlinearities of CIN dendrites. We demonstrate that HCN and NaP currents shape the quasi-linear response properties of CINs, and that dendrites contribute additional phase delays. Furthermore, we show that our analysis can reveal information about dendritic location of membrane nonlinearities, and use the analysis to deduce that both HCN and NaP currents are expressed primarily proximally. Blocking NaP currents pharmacologically revealed that only perisomatic illumination triggers a boosting response. The proximal distribution of NaP currents is further supported by measuring: a) how far autonomously generated backpropagating action potentials (bAPs) actively invade the dendritic arbor; and b) the differential boosting by NaP currents of proximal thalamostriatal vs. distal corticostriatal EPSPs.

## Results

### Ionic currents underlying amplification and resonance in cholinergic interneurons

The HCN and NaP currents depolarize CINs over largely non-overlapping voltage ranges. The HCN current is mostly active below –60 mV and is responsible for the voltage sag in response to a hyperpolarizing current pulse (Figure 1A, “HCN”), whereas the NaP current takes over at –60 mV and is necessary and sufficient (Bennett et al., 2000) to drive CINs to action potential threshold (Figure 1A, “NaP”). Therefore, because NaP is a regenerative current, while HCN is a restorative current, we would expect the current responses to an oscillating voltage command to depend strongly on whether the membrane voltage is clamped above or below –60 mV (Beatty et al., 2015). We therefore held mouse CINs in whole-cell voltage clamp (*n = 10* neurons, *N = 4* mice), first at –55 mV, and subjected them to a voltage command that was composed of a continuous sequence of sinusoidal cycles with an amplitude of 2 mV and a frequency that increases discretely from 0.2 to 20 Hz. The current amplitude was very small at low frequencies and increased monotonically to the high frequency (Figure 1B, black), which is suggestive of an impedance curve with amplified lower frequencies. Loosely speaking, this means that the CINs’ “input resistance” to a low frequency oscillatory input currents is boosted. As expected (Figure 1C, black), the impedance curve, |*Z*(*f*)|, exhibited an amplifying structure, with the phase delay being strictly positive. In order to quantify the degree of amplification, we fit the model of the phase delay, *ϕ*_s_, for an iso-potential cell with quasi-linear properties (see Materials and Methods). In this fit, there is a (negative) amplifying parameter, *µ*_*n*_ – that is derived from biophysical properties of amplifying current (e.g., the slope of the activation curve, reversal potential, etc.) (Goldberg et al., 2007) – which was estimated to be *µ*_*n*_ *= –3.9*.

**Figure 1:**
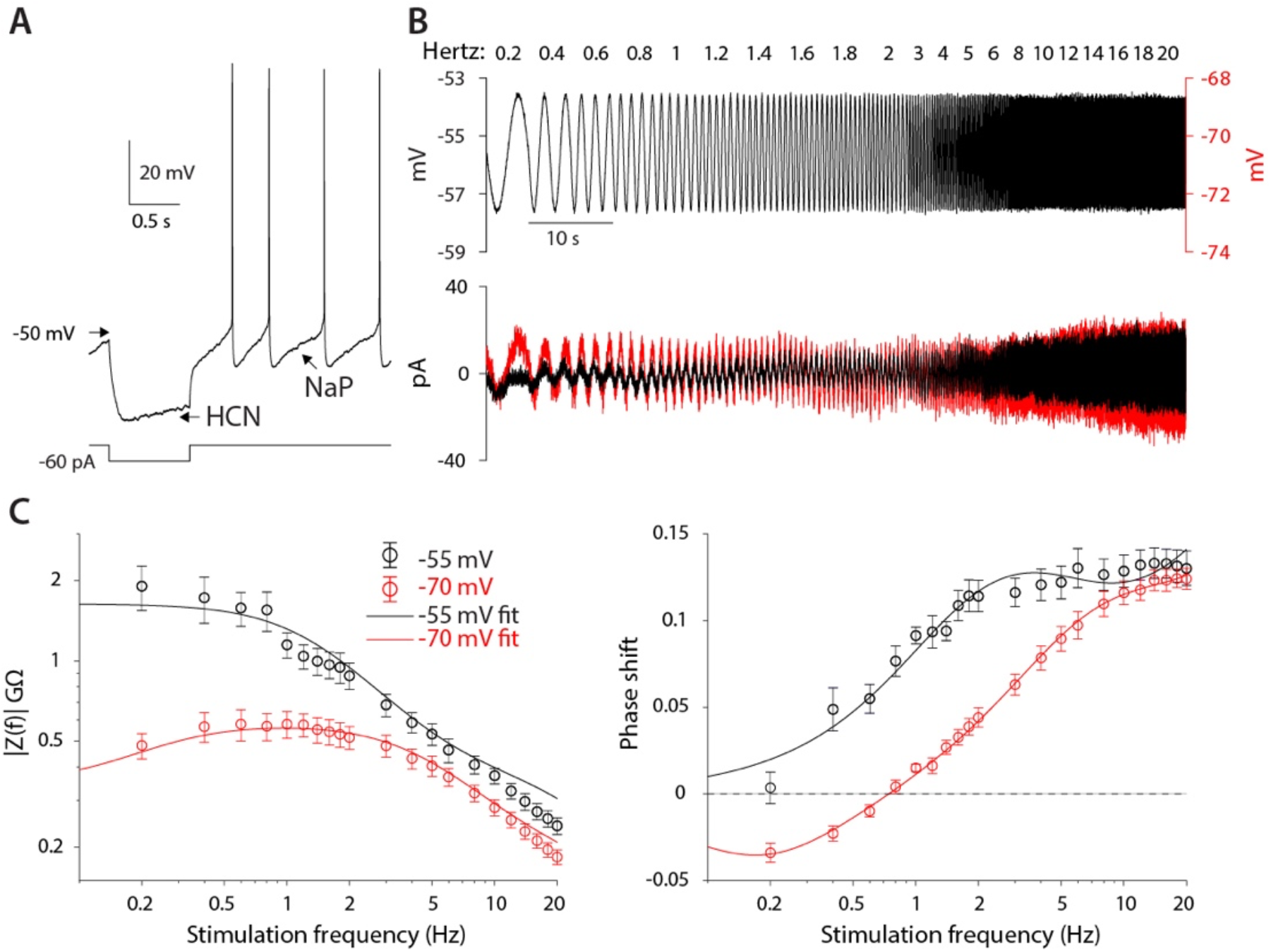
CIN membranes exhibit voltage-dependent quasi-linear properties. **A**. CINs exhibit a voltage sag due to HCN currents, and autonomous pacemaking due to NaP currents. **B**. Application of 2 mV sinusoidal voltage commands to the soma, of increasing frequencies, elicits a current response that is voltage dependent (black trace, –55 mV; red trace, –70 mV). **C**. Estimation of the impedance (left) and phase shift (right) show that at –55 mV, CINs exhibit an amplified impedance and that at –70 mV, CINs exhibit a resonance (non-monotonic impedance and negative phase delays). Solid lines are parameter fits for (α^2^ + β^2^)^-1/2^ shift, *ϕ*_s_ (right, see Eq. 5).

When neurons were held at –70 mV (*n = 10* neurons, *N = 7* mice), the current response was very different, it was much larger at subhertz frequencies as compared to the experiment at –55 mV, and then exhibited what looks like a slightly decreasing amplitude for frequencies near 1 Hz, followed by an amplitude increase at higher frequencies (Figure 1B, red). Estimation of the amplitude response and phase delay (Figure 1C, red), revealed significantly different curves (amplitude: *P = 6·10*^*-25*^, phase: *P = 1·10*^*-17*^, ANCOVA) with a clear resonance peak at approximately 1 Hz, and a zero crossing at a slightly lower frequency (Figure 1C, red). Fitting the somatic phase delay yielded a reasonable fit only when both an amplifying and resonant component were included in the fit with the amplifying parameter (*µ*_*n*_ *= –3.4*) being only slightly reduced relative to the –55 mV fit. In contrast, the additional (positive) resonance parameter, *µ*_*h*_ – which is derived from the biophysical properties of the restorative current – was estimated to be *µ*_*h*_ *= 1.6* (see Materials and Methods) based on the fit to the negative lobe in the phase delay.

The previous experiments suggest that the amplifying effect at –55 mV occurs due to prominence of the amplifying NaP current in that voltage range, whereas the resonance visible at –70 mV is due to the HCN current that dominates that voltage range. This conclusion was supported by the fact that application of 1 µM TTX (*n = 9* neurons, *N = 4* mice), that abolished autonomous spiking and slightly hyperpolarized the CIN (Figure 2A), reduced the impedance (*P = 9·10*^*-9*^) and exhibited a trend toward a shortened phase delay (*P = 0.137*, ANCOVA, Figure 2B). These changes are captured by the amplification parameter being estimated as less negative (*µ*_*n*_ *= –2.0*), which reduces *ϕ*_s_ (Eq. 5 in Materials and Methods). Similarly, application of 10 µM ZD7288, the selective HCN antagonist (*n = 9* neurons, *N = 7* mice), which abolished the sag response (Figure 2C), abolished the resonance peak in the impedance curve (*P = 4·10*^*-16*^) and significantly reduced the negative lobe in the phase delay (*P = 0.018*, ANCOVA, Figure 2D), which was captured by the resonance parameter being reduced to *µ*_*h*_ *= 0.5*.

**Figure 2:**
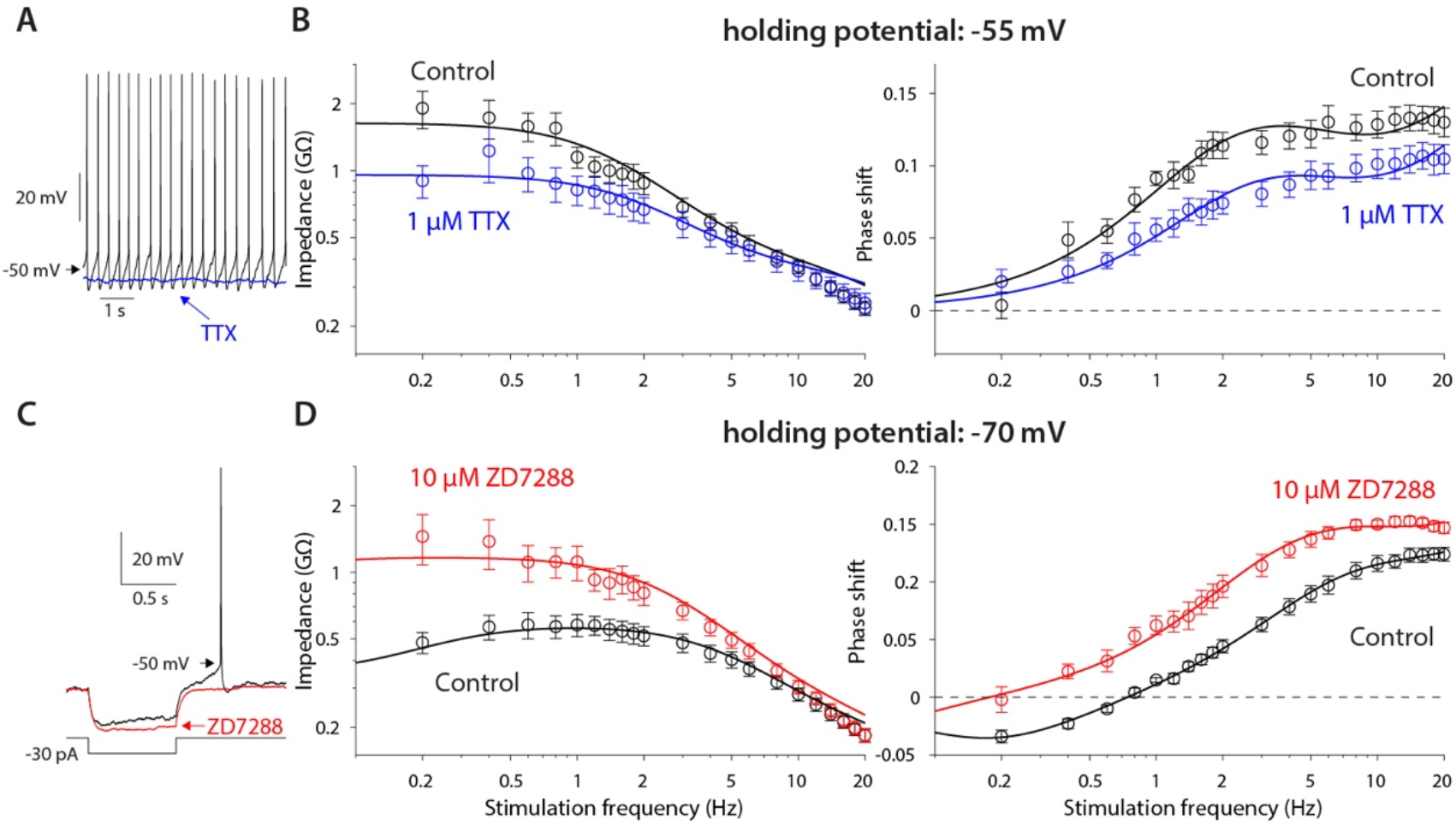
Amplification is caused by NaP currents, whereas the resonance is caused by HCN currents. **A**. TTX (blue) prevents autonomous spiking (black). **B**. TTX prevents amplification of the impedance (left), and reduces the phase shift (right). **C**. ZD7288 (red) abolishes the voltage sag (black). **D**. ZD7288 abolishes the resonance peak in the impedance, and the negative phase shifts in the subhertz range. Solid lines are parameter fits as in Figure 1.

### Optogenetic interrogation of the spatial distribution of NaP and HCN currents in the CIN dendritic arbor

While the above experiments demonstrate the NaP and HCN currents are capable of transforming subthreshold voltage fluctuations, the question remains as to where within the CIN’s somatodendritic compartments these currents perform their amplifying and restorative actions, respectively. One extreme scenario is that they are restricted to the axosomatic region (where they are needed for sustaining the autonomous firing patterns of the CINs). In that scenario, the dendrites could be entirely linear, passively transmitting the distal depolarizations to the soma. Only at the soma are the inputs then transformed by these currents. However, a more realistic scenario is that these currents are also expressed dendritically and exert their nonlinear influence on synaptic inputs more distally. But in this scenario, we would still want to know where along the dendrites the currents are expressed.

To determine the precise location of these channels would require advanced *in situ* molecular and anatomical techniques and/or direct electrophysiological recordings from CIN dendrites in conjunction with advanced imaging techniques. While these experiments may, in principle, be done, we wondered whether the optogenetics-based technique that we recently developed to interrogate the role of dendrites in synaptic integration (Tiroshi and Goldberg, 2019) could help us address this question.

We previously showed that the impact of dendrites on synaptic integration can be quantified by studying the response of neurons that express ChR2 post-synaptically to (470 nm) illumination of their dendritic arbor. In particular, we compared between two spatial patterns of illumination of the somatodendritc arbor of SNr neurons: either a small perisomatic region (approximately 100 µm in diameter) or the entire dendritic arbor. By using sinusoidally-modulated blue LED illumination at various temporal frequencies, we were able to calculate the phase delays produced by both spatial patterns, and found that illumination of the entire dendritic arbor introduced larger phase delays. In order to quantify the effect, we fit the data to a tractable theoretical model of a semi-infinite cable (Goldberg et al., 2007; Tiroshi and Goldberg, 2019). As mentioned above, in the case of the SNr neuron, we found that the dendrites were well-fit by a passive linear cable model, whose parameters (i.e., time and space constants) we could estimate. The conclusion that SNr dendrites were largely linear was further supported by the finding that these phase delays were voltage-independent (Tiroshi and Goldberg, 2019).

Because CINs exhibit prominent amplifying and resonating currents that are strongly voltage dependent (Figures 1&2), we posited that by using the same optogenetic technique and semi-infinite cable model we would be able to quantify the contribution of CIN dendrites to post-synaptic integration. In particular, we hypothesized that by fitting a *quasi-linear* cable model, we would be able to quantify to what degree CIN dendrites *per se* possess amplifying or resonating properties (See Materials and Methods). Finally, by comparing illumination of the proximal *vs*. the entire dendritic arbor, we could learn something about the localization of the nonlinearities along the dendritic arbor. To this end, we crossed mice that express Cre-recombinase under a ChAT promoter with the Ai32 mouse that expresses ChR2 and EYFP in a Cre-dependent manner (see Materials and Methods). The cholinergic neuropil and individual CINs could be clearly visualized in the dorsal striatum of these ChAT-ChR2 mice (Figure 3A). Individual CINs were patched and recorded in voltage clamp, while illuminating either the proximal region with a 60X water-immersion objective (Figure 3A) or the entire slice with a 5X air objective, with a continuous sequence of sinusoidally modulated illumination waveforms at various frequencies (Figure 3B, blue).

**Figure 3:**
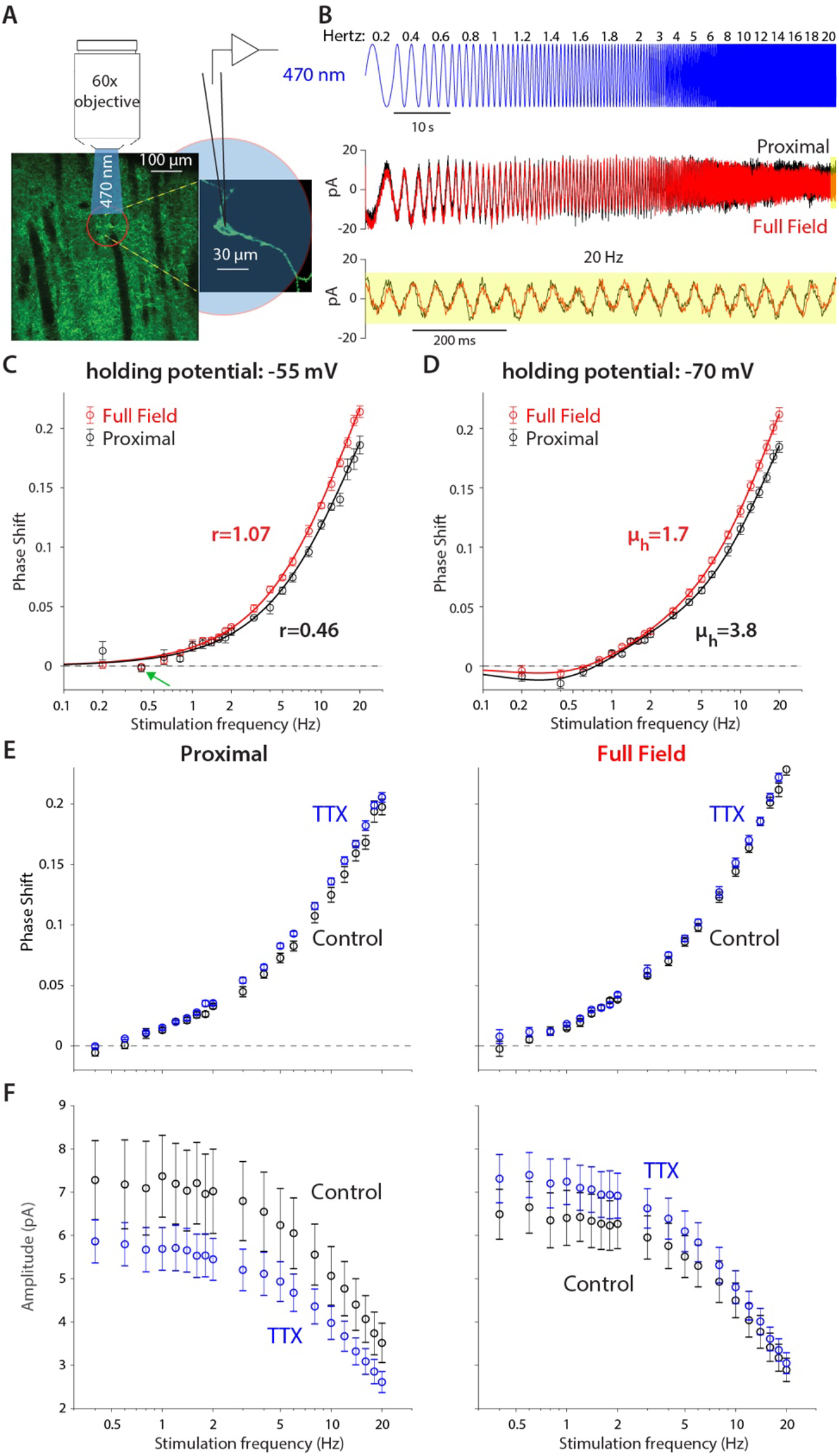
Optogenetic interrogation of the quasi-linear properties of CIN dendrites indicates that dendritic nonlinearities are more prominent proximally. **A**. A CIN in a sagittal slice from ChAT-ChR2 mouse is patch-clamped in the whole-cell mode while either a small proximal region around the soma or the full-field are illuminated with a sinusoidally modulated 470 nm LED. **B**. The current response to the proximal (black) and full-field (red) illumination differ, with the phase of the full-field illumination right-shifted at the higher frequencies (20 Hz is highlighted in yellow). **C**. Phase shifts at –55 mV holding potential, calculated for proximal (black) and full-field illumination (red). A tendency towards negative phase shifts is present at 0.4 Hz (green arrow). Fitting the passive model at –55 mV demonstrated that the effective range of illumination (*r*) is larger for the full field fit (Eq. 2). **D**. Phase shifts at –70 mV holding potential, exhibit a negative phase shift, and the resonance parameter (*µ*_*h*_) is smaller for the full field fit, as is the magnitude of the amplification parameter (*µ*_*n*_, see main text). The elevation in these parameters’ magnitudes when illuminating proximally relative to full-field suggest that the surface densities of NaP and HCN currents is higher proximally. **E**. Phase response for proximal (left) and full-field illumination (right) in TTX at –55 mV. **F**. Amplitude response for proximal (left) and full-field illumination (right) at –55 mV reveals an opposite effect of TTX.

Comparison of the somatic current traces in response to proximal (Figure 3B, black) *vs*. full-field (red) illumination demonstrated that the phase of the full-field-generated current is delayed relative to the proximally-generated current. This could be observed in the raw data both for the low and high (yellow inset) frequencies. The effect at the high frequency was very evident when plotting the phase delay curves both when the CINs were held at –55 mV (Figure 3C) and at –70 mV (Figure 3D). Estimation of these phase delays revealed delays that were considerably larger than those observed with the electrical somatic stimulation (Figure 1). The main contributor to the large delays are the kinetics of the ChR2, *ϕ*_c_ (Eq. 6b, in Materials and Methods) with an additional dendritic delay *ϕ*_d_ (Eq. 2 in Materials and Methods). So we used independent measurements of the ChR2 kinetics in CINs (Figure 3–figure supplement 1), and previous literature about these kinetics (Nagel et al., 2003; Tchumatchenko et al., 2013) to fit the phase and amplitude contribution of ChR2, as explained in Appendix 1.

We found that fitting our model of ChR2 kinetics to the significantly-different phases observed in the proximal and full-field illuminations at – 55 mV (*n = 5* neurons, *N = 4* mice, *P = 4·10*^*-3*^, ANCOVA) yielded that the full-field illumination activated roughly twice the electrotonic range (*r = 1.07*) that was activated by the proximal illumination (*r = 0.46*, Figure 3C). However, a closer look at these fits reveals that the phase delay at 0.4 Hz (for both the proximal and full field illumination), is not captured by this model. This dip in phase delay (Figure 3C, green arrow) at this low frequency is reminiscent of what a restorative current is expected to do. In order to accentuate the effect of the restorative HCN currents, we repeated the experiment at the –70 mV holding potential (Figure 3D). In this case (*n = 14* neurons, *N = 10* mice), the phase delays at the lower frequencies (especially at 0.4 Hz) were negative, which was reminiscent of the results from the electrical voltage stimulation experiments (Figure 1) at the –70 mV holding potential. Estimation of the amplitude responses for proximal and distal stimulation at both holding potentials revealed that they were less sensitive at revealing the resonance structure (Figure 3–figure supplement 2), that was more readily read off from the phase responses (Figure 3D) in the sense that phase estimates provided tighter error bars than the amplitude estimates.

When fitting a full ChR2 kinetics plus quasi-linear dendrite model (incorporating both amplifying and restorative parameters) to the –70 mV measurements (Figure 3D), we found that the curves were significantly different (*P = 8·10*^*-4*^, ANCOVA) and the estimates of the effective electrotonic range of illumination remained similar to those estimate with the passive model (Figure 3C): *r = 1.06* for the full-field and *r = 0.58* for the proximal illumination. Here too the restorative parameter was estimated to be larger for the proximal fit (*µ*_*h*_ *= 3.4*) relative to the full field fit (*µ*_*h*_ *= 1.7*), suggesting that the HCN currents are denser more proximally.

Table 1 summarizes the model parameters used to fit the curves in Figures 1-3. In Appendix 1, we discuss how the various parameters affect the model and provide a more detailed description of how the parameter space was searched to fit the model.

**Table 1.** Parameters fit to quasilinear model in Figures 1–3.

| Figure, Curve | $\mu_n$ | $\mu_h$ | $\tau_n$ (ms) | $\tau_h$ (ms) | $\gamma_R$ | $\tau$ (ms) | $r$ | Amp (GΩ) |
| --- | --- | --- | --- | --- | --- | --- | --- | --- |
| 1C, -55 mV | -3.9189 | - | 28 | - | 5.4296 | 41.8 | - | 2.4728 |
| 1C, -70 mV | -3.3854 | 1.5863 | 11.6 | 1077.8 | 5.8722 | 26.1 | - | 1.4229 |
| 2B, TTX | -2.0178 | - | 31.7 | - | 3.4610 | 19.4 | - | 1.3809 |
| 2B, ZD7288 | -1.8988 | 0.4844 | 13.4 | 9107.5 | 2.6111 | 17.2 | - | 0.8323 |
| 3C, proximal | - | - | - | - | - | 35.8 | 0.4360 | - |
| 3C, distal | - | - | - | - | - | 26.8 | 1.0651 | - |
| 3D, proximal | -1.4763 | 3.4400 | 60.0 | 551.2 | 3.3405 | 33.6 | 0.5788 | - |
| 3D, distal | -0.8885 | 1.6646 | 49.2 | 403.3 | 3.8850 | 45.3 | 1.0646 | - |

The conclusion that the HCN current is concentrated proximally is buttressed by the fact that the negative phase delays in the subhertz frequency range tended to be less negative for the full-field illumination (*P = 0.037*, Wilcoxon rank sum test on phase delays at 0.4 Hz). In the framework of our quasi-linear model, the only mathematically possible way to recreate this finding while increasing the effective electrotonic range *r* being illuminated is by reducing the restorative parameter *µ*_*h*_. Otherwise, if *µ*_*h*_ were constant then illuminating a larger region of the cable – by increasing *r* – would necessarily recruit more restorative current, thereby making the negative delays more negative (See Appendix 1).

Aside from the deviation at 0.4 Hz (Figure 3C, green arrow, which may be indicative of a resonant component), the other phase measurements at –55 mV are consistent with a simple model of ChR2 kinetics plus a passive dendritic delay, thereby raising the possibility that optogenetic dendritic activation fails to engage NaP currents. This failure could suggest that NaP currents are relatively absent from the dendrites. However, our somatic measurements showed clear evidence that NaP currents affect the quasi-linear properties of the soma (Figure 2B). Additionally, the fit to the phases at –70 mV, required the inclusion of an amplification parameter, which was estimated to have a larger magnitude for the proximal illumination (*µ*_*n*_ *= –1.5*) than for the full-field illumination (*µ*_*n*_ *= –0.9*). It therefore seems more likely that the Nav channels that underlie the NaP currents are localized proximally and taper off distally. To test this, we repeated the measurements at –55 mV before and after application of TTX. While TTX application may have slightly increased the phase delays for both proximal and full-field illumination (Figure 3E) – which could reflect a reduction in the overall dendritic membrane conductance of the CINs, and therefore a lengthening of its space constant – the amplitude responses (Figure 3F) unequivocally show that TTX exerts an *opposite* affect when proximal *vs*. full-field illumination are used. While full-field illumination in TTX increased the somatic current’s amplitude response (control: *n = 9* neurons, *N = 7* mice; TTX: *n = 14* neurons, *N = 8* mice, *P = 7·10*^*-9*^, ANCOVA) presumably by reducing the current escape via the dendritic membrane, proximal illumination in TTX reduced the somatic current’s amplitude response (control: *n = 9* neurons, *N = 7* mice; TTX: *n = 12* neurons, *N = 6* mice, *P = 1.1·10*^*-4*^, ANCOVA) indicating that proximal NaP currents indeed boost the somatic current, as concluded from the somatic experiments (Figure 2). Thus, this pharmacological result provides *model-independent* evidence that the NaP currents are expressed primarily proximally and less so distally. In the following section, we provide independent evidence in support of this conclusion.

### Distance of dendritic bAP invasion indicates location of amplifying Nav channels

The persistent and fast-inactivating Nav currents flow through the same Nav channels (Alzheimer et al., 1993). Therefore, a method that is indicative of where these NaV channels are located will indicate where the NaP current can be found. One such method involves determining to what distance from the soma dendritic bAPs invade the dendritic arbor. To this end, we used 2PLSM Ca^2+^ imaging to measure the size of the Ca^2+^ transients elicited by bAPs in autonomously firing CINs at various distances from the soma (Figure 4A). We conducted line scans (*n = 11* neurons, *N = 7* mice) to measure the Ca^2+^ signals at various distances from the soma (Figure 4B). Next, we estimated the size of the spike triggered average (STA) of the *ΔF/F*_*0*_ Ca^2+^ signal, by averaging around spontaneous APs measured at the soma (Figure 4C). The amplitude of the STA was estimated by fitting it with an alpha-function (see Materials and Methods). A scatter plot of STA amplitude *vs*. distance from soma demonstrates a large degree of variability (Figure 4D). Nevertheless, applying a 35 µm moving average to the scatter plot reveals a trapezoidal dependence of Ca^2+^ transient sizes (Figure 4D, black). We previously demonstrated that somatic Ca^2+^ transients are smaller than dendritic transients due to the difference in the surface-area-to-volume ratio (Goldberg et al., 2009; Rehani et al., 2019). In the same vein, the initial dip in the size of the transients at short distances from the soma result from the large size of the proximal dendrites (Figure 4D), relative to the distal dendrites. Neglecting that effect, we found that the size of the bAP-driven Ca^2+^ transients remains constant up to approximately 70 µm from the soma, and then drops off. Additionally, in 7 CINs (*N = 5* mice) in which we had measurements of *ΔF/F*_*0*_ at both proximal and distal (>70 µm) locations, we found that the median distal signal was significantly lower than the proximal one by 29% (*P = 0.047*, Wilcoxon rank sum test). This spatial dependence of the Ca^2+^ transient amplitude, suggests that the bAP maintains a constant amplitude due to the presence of Nav channels that sustain their regenerative nature out to some 70 µm from the soma. Farther out the Ca^2+^ transients decrease presumably due to a drop off in Nav channel expression, which leads to a lower amplitude bAPs, and hence less Ca^2+^ via voltage-activated Ca^2+^ channels. Thus, this measurement strengthens the conclusion that NaP currents are present in CIN dendrites – primarily in proximal dendrites (up to approximately 70 µm from the soma).

**Figure 4:**
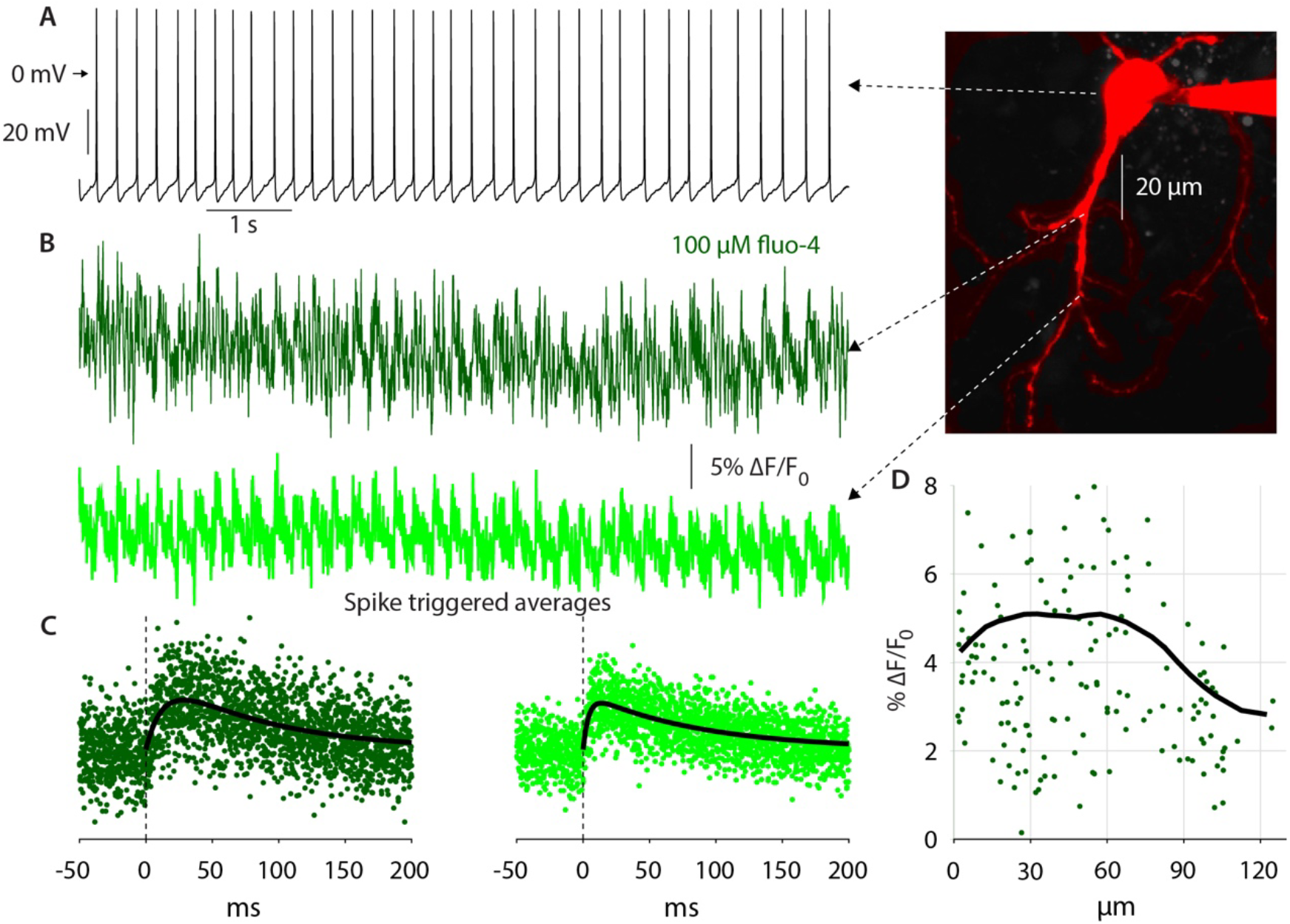
Autonomous action potentials actively back-propagate in CIN dendrites up to 70 µm from the soma. **A**. Autonomous discharge of a CIN that was patched and filled with fluo-4 and Alexa Fluor 568 for 2PLSM imaging (image). **B**. Line scans at various distances from the soma exhibit Ca^2+^ oscillations caused by bAPs. **C**. Calculating the spike-triggered average of these oscillations and fitting an alpha-function gives an estimate of the amplitude of these oscillations (in % *ΔF/F*_*o*_). **D**. The scatter plot of these amplitudes as a function of distance from the soma (11 CINs from 7 mice are pooled) exhibits a large degree of variability. However, a 35 µm moving average (black line) exhibits that the Ca^2+^ transients begin to decay approximately 70 µm from the soma, indicating that bAPs are supported by Nav channels up to that point (the initial increase up to 20 µm is due to the increase in the surface-to-volume ratio in the large proximal dendrites).

### NaP currents boost proximal thalamic inputs but not distal cortical inputs to CINs

The above data indicate that Nav channels are preferentially expressed in proximal dendrites. These channels should amplify synaptic inputs impinging on the dendrites. In the case of linear dendrites – either passive or even quasi-linear dendrites – the relative boosting will affect all inputs at all locations equally. However, as demonstrated in Appendix 2 – using a simulated nonlinear dendrite that expresses the NaP current proximally – proximal inputs are more strongly *nonlinearly* boosted than distal inputs, because the latter decay passively before reaching the amplifying region, and are therefore boosted less.. Thus, because thalamic inputs on to CINs terminate proximally, whereas cortical inputs terminate distally (Doig et al., 2014; Lapper and Bolam, 1992; Mamaligas et al., 2019; Thomas et al., 2000), in order to generate comparable EPSPs we presumably need to activate many more distal cortical inputs than proximal thalamic inputs. However, because distal cortical inputs are presumably amplified less their spatial summation at the soma will also be amplified less. In contrast, because proximal thalamic inputs consist of individual inputs, each of which engages the local nonlinear amplification, their summation at the soma will also exhibit more amplification. Thus, we expect only the thalamic – but not cortical – inputs to be significantly boosted by Nav current. To test this, we transcranially inoculated the parafascicular nucleus (PfN) of Vglut2-ires-Cre mice with adeno-associated viruses (AAVs) harboring Cre-dependent ChR2 and EYFP genes (Figure 5A) (Aceves Buendia et al., 2019; Rehani et al., 2019). Two weeks later, we tested the sensitivity of optogenetically-evoked monosynaptic EPSPs (Figure 5–figure supplement 1A) to ranolazine (30 µM), a selective antagonist of the NaP current, in CINs patched in acute striatal slices. The mean amplitude of the EPSP was reduced by a median of 16% (*n = 11* neurons, *N = 4* mice, *P = 0.04*, Wilcoxon signed-rank test, Figure 5B). In order to test whether the more distal cortical inputs are modulated by NaP, we used the Thy1-ChR2 mice (Aceves Buendia et al., 2019; Arenkiel et al., 2007) that expresses ChR2 in corticostriatal fibers (Figure 5C), but neither in the PfN (Figure 5–figure supplement 1B) nor in the pedunculopontine nucleus (Gradinaru et al., 2009) that also provides monosynaptic glutamatergic projection to CINs (Assous et al., 2019). The monosynaptic EPSPs (Figure 5–figure supplement 1A) elicited by cortical inputs to CINs in these mice were unaffected by ranolazine (*n = 11* neurons, *N = 6* mice, *P = 0.74*, Wilcoxon signed-rank test, Figure 5D). Moreover, ranolazine had no effect on paired-pulse ratios (Aceves Buendia et al., 2019) at either synapse (Figure 5–figure supplement 1C), ruling out a presynaptic mechanism-of-action. Thus, we conclude that the post-synaptic modulation of glutamatergic inputs to CINs by NaP currents corresponds to the spatial distribution of these currents as derived from the quasi-linear properties of CINs and from our 2PLSM imaging of bAPs.

**Figure 5:**
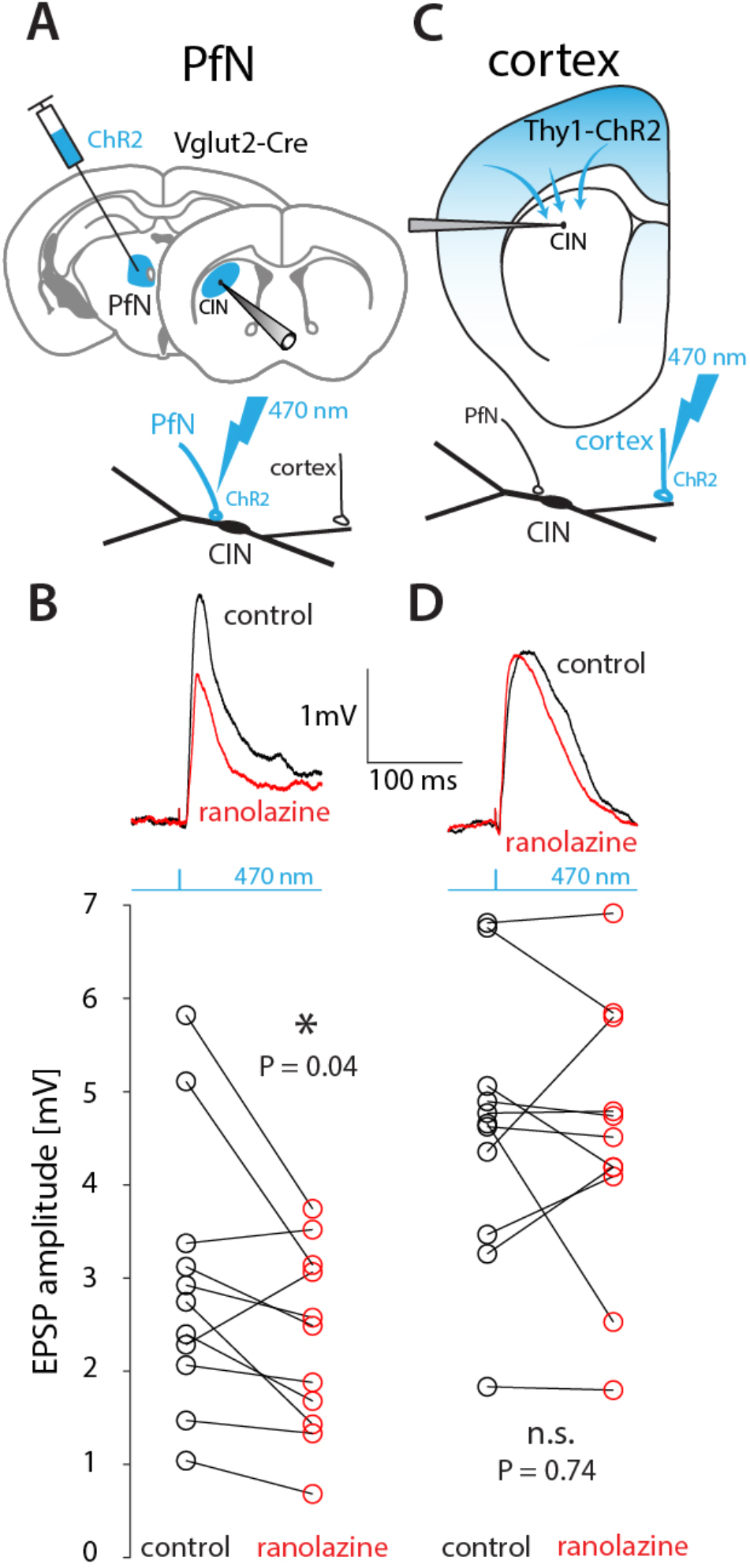
Thalamic – but not cortical EPSPs – onto CINs are boosted by NaP currents in healthy mice. **A**. The parafascicular nucleus (PfN) of Vglut2-Cre mice was inoculated with AAVs harboring Cre-dependent ChR2, so that 470 nm LED illumination of striatal slices activated monosynaptic PfN terminals while CINs were patched in current clamp mode. **B**. Optogenetically evoked thalamic EPSPs in CINs (held between –50 mV and –60 mV) were reduced by 30 µM ranolazine. **C**. CINs were patched in current clamp in Thy1-ChR2 mice, so that 470 nm LED illumination of striatal slices activated nominally cortical terminals. **D**. Optogenetically evoked monosynaptic cortical EPSPs were unaffected by ranolazine.

### Tonically active neurons in non-human primates exhibit a pause-like response to slow wave oscillations during sleep

Our data, alongside previous studies (Beatty et al., 2015), demonstrate that somatodendritic HCN currents give rise to resonances in the subhertz range in CIN surface membranes. In contrast, SPNs exhibit no resonance (Beatty et al., 2015). This suggests CINs in intact animals may exhibit increased sensitivity to oscillatory inputs in that frequency range, whereas SPNs should not. Delta waves in the electroencephalogram (EEG) and in cortical and sub-cortical local field potentials (LFP) are prominent during non-REM sleep across species (Brown et al., 2012; Liu and Dan, 2019), including in non-human primates (NHPs) (Mizrahi-Kliger et al., 2018). Because the LFP is widely thought to represent subthreshold cellular activity, affected by afferent and recurrent synaptic inputs (Buzsáki et al., 2012), we hypothesized that the tonically active neurons (TANs) in the NHP striatum – that are comprised primarily of CINs (Aosaki et al., 1995; Kawaguchi, 1993; Wilson et al., 1990) – will exhibit stronger entrainment to slow-wave LFP events during sleep, while SPNs under the same conditions will not. Indeed, triggering the spike trains of TANs and SPNs (from *N = 2* NHPs) on slow wave events that occur during sleep stages N2 and N3 (Mizrahi-Kliger et al., 2018; Riedner et al., 2007), which represent relatively deep non-REM sleep, demonstrates that while the firing rate of SPNs was unaffected (*n = 83*), the firing rates of TANs (*n = 122*) was modulated by these slow-wave events (Figure 6A-E). In contrast, the TANs firing was not modulated during higher frequency sleep spindle events (Figure 6F-H), in agreement with the subhertz membrane resonance peak we observed in the acute slice experiments. Importantly, the biphasic response exhibited by TANs to slow-wave events bears a striking resemblance to the “classical” TAN response to external cues (Aosaki et al., 1995; Apicella et al., 1991; Joshua et al., 2008; Kimura et al., 1984). Because these responses in awake primates require an intact thalamic projection (Bradfield et al., 2013; Goldberg and Reynolds, 2011; Matsumoto et al., 2001; Schulz and Reynolds, 2013; Smith et al., 2004; Yamanaka et al., 2018) (and there is no reason to assume that this is altered in sleep), our findings (Figure 6) provide support to the hypothesis that the CIN membrane resonance contributes to a thalamically-driven biphasic CIN response during slow-wave sleep, possibly because this projection terminates proximally on CINs. Importantly, because the primates are asleep in a sound-proof room, this is, to the best of our knowledge, the first report of striatal TANs responding to an internally-generated brain event, and not an external saliency-related cue.

**Figure 6:**
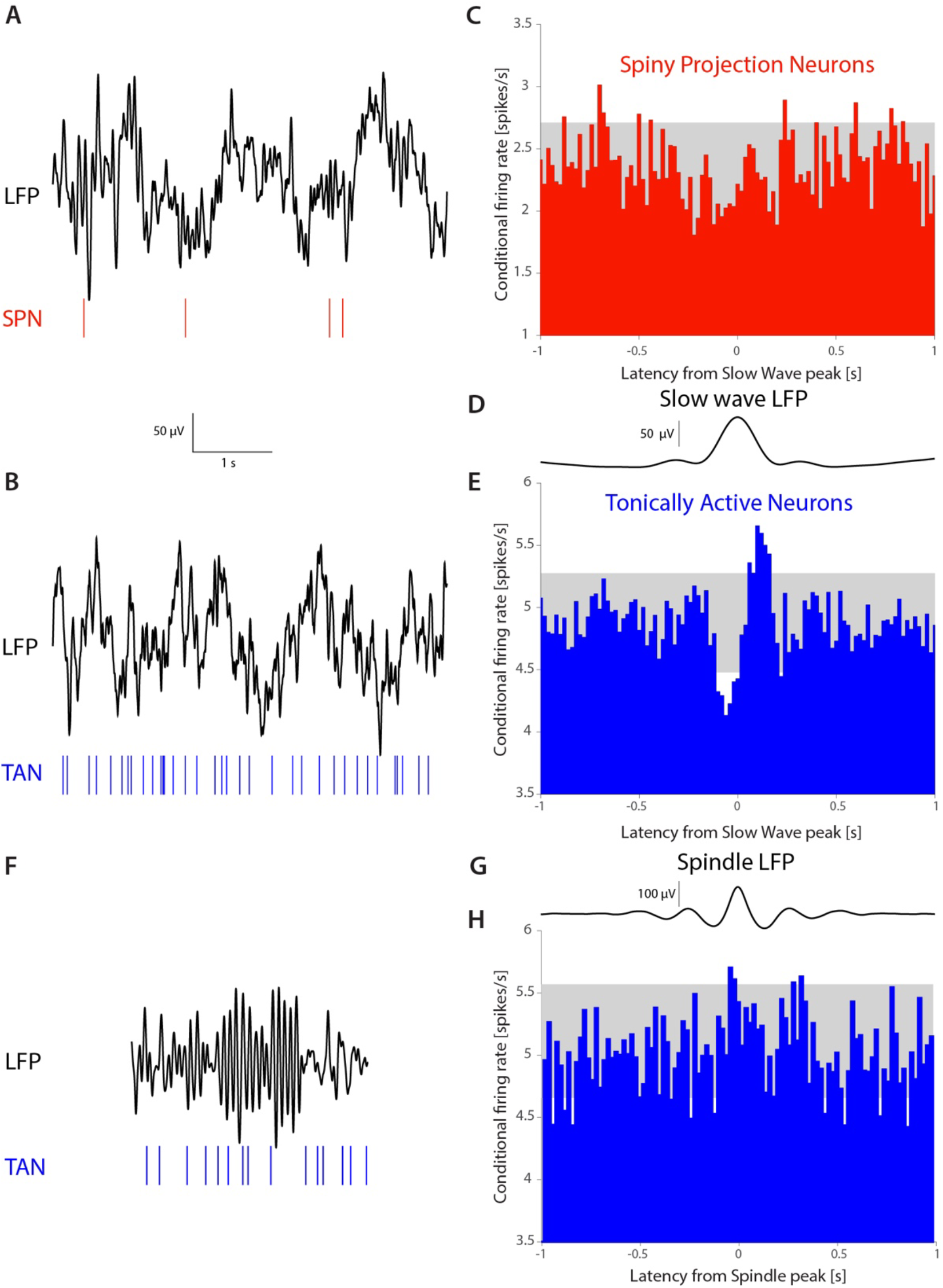
TANs, but not SPNs, exhibit a pause-like response to slow-wave events occurring during natural non-REM sleep in non-human primates (NHPs). **A**. simultaneous recording of LFP and an SPN in an NHP during N2 and N3 stages of sleep. **B**. simultaneous recording of LFP and a TAN in an NHP during N2 and N3 stages of sleep. **C**. SPN firing rate conditioned on the occurrence of an LFP slow wave event (6,065 triggers). **D**. Average striatal LFP signal triggered on the occurrence of slow wave events (see Materials and Methods). **E**. TAN firing rate conditioned on the occurrence of slow wave event (28,603 triggers). **F**. simultaneous recording of LFP and a TAN in an NHP during a sleep spindle. **G**. Average striatal LFP signal triggered on the occurrence of a sleep spindle (see Materials and Methods). **H**. TAN firing rate conditioned on the occurrence of a sleep spindle (5,829 triggers). Gray box indicates the 99% confidence intervals.

## Discussion

### Quasi-linear dendritic properties of CINs and the non-uniform distribution of their active conductances

In the current study, we applied a cable-theoretic and optogenetics-based formalism – that we developed previously to study dendritic properties of SNr neurons (Tiroshi and Goldberg, 2019) – to study the dendritic properties of CINs. Unlike SNr neurons that were found to act like passive linear cables exhibiting no voltage dependent properties, we find that CIN dendrites exhibit both amplification and resonances in a voltage dependent manner: amplification is more prominent at depolarized subthreshold potentials (approximately –55 mV), whereas resonances are more prominent at more hyperpolarized potentials (approximately –70 mV). Moreover, we found that our method is able to provide information about where in the dendritic arbor these quasi-linear properties are localized. We extracted boosting (*µ*_*n*_) and resonance (*µ*_*h*_) parameters for proximal vs. full-field illumination, and found that the magnitude of both parameters is smaller when the entire dendritic arbor is illuminated. In other words, activating more of the distal dendritic membrane dilutes the amplifying and resonating effects, meaning that the additional membrane illuminated in the full-field condition contributed less to amplification and resonances. The simplest interpretation of this finding is that the active conductances that give rise to amplification and resonance are more highly expressed proximally and apparently taper off farther out in the dendritic arbor. We identified the relevant currents by studying the quasi-linear properties of CIN somata with voltage perturbations. We found that amplification is TTX-sensitive and therefore arises from Nav channels, that give rise to the NaP current. Similarly, we found that resonances require HCN channels.

We used one-direct and two indirect methods to validate the preferentially proximal localization of NaP currents. First, TTX application revealed that boosting occurs only at the soma (Figure 2B) and in proximal (but not distal, Figure 3F) dendritic membranes. Importantly, the TTX results demonstrate that our conclusion that boosting is restricted to the soma and proximal dendrites is *independent* of the quasi-linear model fit. Second, we implemented a widely-used approach of estimating the distance of dendritic invasion of bAPs by imaging the Ca^2+^ influx that accompanies them (Carter and Sabatini, 2004; Day et al., 2008; Goldberg et al., 2009; Kerr, 2004; Rehani et al., 2019; Tanimura et al., 2016). We found – not unlike in SPN neurons – that bAPs maintain their amplitude out to 70 µm from the soma and then begin to decay, suggesting that Nav currents that support bAP propagation taper off from that point onwards. Because strictly speaking a change in Ca^2+^ transients could result from a change in the concentration of voltage-activated Ca^2+^ channels, future work can verify our results by directly testing where locally puffed TTX attenuates the bAP (perhaps even with the use of voltage sensitive probes). Second, we demonstrated that only thalamic – but not cortical – EPSPs exhibit sensitivity to the selective NaP blocker. Because the thalamic terminals are located more proximally (Lapper and Bolam, 1992; Mamaligas et al., 2019; Thomas et al., 2000), this provides further evidence that NaP currents are more prominent locally. The preferential localization of HCN currents proximally in CIN dendrites requires additional validation, particularly because other neurons exhibit an opposite pattern (Berger et al., 2001; Harnett et al., 2015; Kole et al., 2006). However, we were less interested in validating the proximal localization of HCN currents, because unlike NaP currents, activation of HCNs is conditioned on CINs being actively (after-)hyperpolarized, during which bAPs are unlikely to occur. We did however test a context in which HCN currents, and the resonance they underlie, are likely to affect CIN collective dynamics. We found that TANs respond biphasically to internally-generated slow wave activity during sleep, but not to sleep spindles. This preferential entrainment is consistent with the fact that CIN membranes exhibit a resonance in the subhertz range as found by us and others (Beatty et al., 2015). The elevated impedance (or frequency-dependent access resistance) in this range means that inputs fluctuating in this range will more efficiently depolarize the CINs, and therefore more likely to trigger additional action potentials than inputs fluctuating at a frequency that is far from the resonance frequency. Accordingly, SPNs that do not exhibit resonances (Beatty et al., 2015) are not entrained by the slow wave oscillation. Our findings in sleeping NHPs are at odds with various rodent studies that found that cortical slow-wave activity is weakly associated with the discharge of TANs in anesthetized rodents, and more strongly associated with SPN activity (Goldberg et al., 2003; Mahon, 2001; Reig and Silberberg, 2014; Schulz et al., 2011; Sharott et al., 2012; Stern et al., 1998). These differences may be attributable to species differences and/or differences between natural sleep and anesthesia, which may differ in the degree to which thalamic *vs*. cortical inputs engage striatal neurons. Moreover, the dendritic nonlinearities in SPNs (Plotkin et al., 2011) may be preferentially engaged during anesthesia, thereby causing SPNs to respond more strongly under anesthesia than during natural sleep.

### Advantages and limitations of the quasi-linear formalism and the optogenetics-based approach

As mentioned in the Introduction, the quasi-linear approach provides a tool to characterize membrane excitability in general functional terms, without requiring the complete biophysical characterization of membrane currents. Unfortunately, the method is not invertible: even after estimating all the quasi-linear parameters one cannot deduce the full biophysical characterization of the underlying channels even if one has a precise mathematical model of these channels. Moreover, even though we extracted the values of *µ*_*n*_ and *µ*_*h*_, we do not know how to translate them into a quantitative measure of channel density. Still, the ability to compare parameters for various spatial illumination patterns enables us to reach qualitative conclusions regarding relative channel density. Additionally, our attribution of *µ*_*n*_ to NaP and *µ*_*h*_ to HCN is only provisional. CINs possess other channels that have a major influence on their firing patterns such as A-type, inward rectifying, and various Ca^2+^-activated K^+^ channels, to mention a few (Bennett et al., 2000; Deng et al., 2007; Goldberg et al., 2009; Goldberg and Wilson, 2010, 2005; McGuirt et al., 2021; Oswald et al., 2009; Song and Surmeier, 1996; Wilson, 2005; Wilson and Goldberg, 2006), that were not included in our analysis. Future work should elaborate how these and other channels contribute to the quasi-linear properties of CINs and their dendrites.

Using optogenetics provides another level of practicality. The seminal studies that characterized the filtering properties of dendrites or pyramidal neurons (Berger et al., 2001; Goldberg et al., 2007; Hutcheon et al., 1996; Ulrich, 2002) required dual soma-dendrite patching which is not practical for all neuronal types whose dendrites taper off rapidly. Expressing opsins in the membrane being activated, as we did, means that a single somatic patch electrode can suffice to conduct the quasi-linear characterization. We currently use this approach to illuminate large regions of the dendritic arbor simultaneously, which means we can only derive large-scale dendritic properties. In the future, localized laser stimulation of dendritic regions (visualized with the help of a fluorescent marker in the patch pipette), or even two-photon laser activation of opsins (provided one can find stimulation parameters that do not harm the dendrites) could be used as an alternative approach. On the backdrop of optogenetics being utilized almost exclusively to study circuit mapping (Häusser, 2021; Kim et al., 2017; Petreanu et al., 2009) by expressing opsins presynaptically, we believe our study joins other studies (Higgs and Wilson, 2017; Tiroshi and Goldberg, 2019) to underscore the value of using opsins expressed postsynaptically to study neuronal – even dendritic – excitability.

### Somatic amplification and resonance in CINs

We preceded our optogenetic characterization of dendritic nonlinearities with the electrical characterization of quasi-linear somatic properties using a sinusoidal voltage command applied through the patch pipette. As previously reported by Beatty and collaborators (Beatty et al., 2015), CIN somata exhibit a membrane resonance in the subhertz range, that depends on the holding potential. However, while Beatty and collaborators found that an amplified resonance is present in the more depolarized (approximately – 55 mV) range, we found that the resonance is only pronounced in the more hyperpolarized (approximately –70 mV) range, where it depends on the HCN current. It is possible that the resonances observed by Beatty and collaborators (Beatty et al., 2015) at –55 mV arose from ineffective clamping of dendrites with the slightly higher resistance electrodes used in that study. In support of this proposition, we also found evidence for the resonance arising from unclamped (dendritic) membranes. When the CINs were clamped at –55 mV, while somatic voltage perturbations did not produce a resonance, optogenetic stimulation (both proximal and full field) at 0.4 Hz produced a more negative phase than at the neighboring 0.2 Hz and 0.6 Hz stimulation (Figure 3C, green arrow). This downward deflection in phase is reminiscent of the fullblown negative subhertz region that occurred with optogenetic stimulation when the CINs were clamped at –70 mV (Figure 3D). Thus, it is likely that resonance originated in both studies from dendritic HCN currents. Because in the study of Beatty and collaborators (Beatty et al., 2015) TTX lowered the impedance, without removing the subhertz resonance peak, we conclude that the effect of TTX in both studies is to remove the NaP-dependent boosting, without directly affecting resonance. It also seems that the location of the peak amplitudes estimated by Beatty and collaborators (Beatty et al., 2015) is perhaps 1 Hz larger than the location of the peaks in our study. However, this difference could result from the fact that they used a chirp stimulation where the frequency increases continuously, whereas we used several periods of perfect sinusoidal waveforms with a discrete set of frequencies. Moreover, it can be shown mathematically that for the quasi-linear model the zero crossing in the phase response occurs at a slightly smaller frequency than the peak in the amplitude response, so this may also contribute to the impression that the resonance frequency observed in our experiments is slightly different from that observed in Beatty and collaborators (Beatty et al., 2015).

### Dendritic contribution to capacity of CINs to differentiate between their excitatory inputs

In NHPs, TANs that are comprised primarily of CINs (Aosaki et al., 1995; Goldberg and Wilson, 2010; Kawaguchi, 1993; Wilson et al., 1990), encode – through a brief pause in their tonic firing – external stimuli that are salient (and often unexpected, even conveying a stop or behavioral shift signal (Aoki et al., 2018; Thorn and Graybiel, 2010)) or stimuli that are associated with reward (Apicella et al., 1991; Goldberg and Reynolds, 2011; Kimura et al., 1984). The pause response requires an intact projection from the PfN of the thalamus (Bradfield et al., 2013; Goldberg and Reynolds, 2011; Matsumoto et al., 2001; Schulz et al., 2011; Schulz and Reynolds, 2013; Smith et al., 2004; Yamanaka et al., 2018), indicating that TANs are attuned to thalamic input. While TANs do not respond to ongoing movement (Aosaki et al., 1994; Apicella et al., 1991; Kimura et al., 1984; Raz et al., 1996), and hence seem less attuned to sensorimotor cortical input (Sharott et al., 2012), CIN activity in awake behaving mice is strongly modulated by self-initiated movements (Gritton et al., 2019; Howe et al., 2019; Yarom and Cohen, 2011). Nevertheless, slice physiology studies in rodents (Aceves Buendia et al., 2019; Assous, 2021; Assous et al., 2017; Ding et al., 2010; Johansson and Silberberg, 2020; Kosillo et al., 2016; Threlfell et al., 2012) clearly show that thalamic inputs to CINs are stronger in the sense that they give rise to larger EPSPs (Johansson and Silberberg, 2020) and can trigger a pause-like responses (Ding et al., 2010), whereas cortical inputs to most CINs (Mamaligas et al., 2019) are weaker in that they give rise, in acute slices, to smaller EPSPs and cannot trigger the pause-like response – although this distinction is less pronounced in anesthetized rodents (Doig et al., 2014). Thus, it is clear that CINs differentiate between these two excitatory inputs and respond differently to them. A major contributor to the CINs’ capacity to dissociate thalamic and cortical input is the differential distribution of their respective terminals on the dendritic arbor: thalamic input terminate perisomatically and on proximal dendrites, whereas cortical input terminates on distal dendrites (Doig et al., 2014; Lapper and Bolam, 1992; Mamaligas et al., 2019; Thomas et al., 2000). Our finding that the higher expression of NaP currents in proximal dendrites preferentially boosts thalamic over cortical inputs, suggests that active conductances are expressed in CIN dendrites in a manner that corresponds and reinforces the effect of the spatial separation between the terminals of the two inputs.

### Adaptive changes in dendritic excitability in movement disorders

The capacity of the CIN dendritic excitability to mirror the distribution of afferent terminals is also an adaptive process. We previously reported that CIN dendritic excitability is elevated in the Q175 mouse model of Huntington’s disease (HD) (Tanimura et al., 2016). In contrast to control (Thy1-ChR2) mice in which cortical EPSPs in CINs are insensitive to ranolazine (Figure 5), cortical excitatory postsynaptic currents (EPSCs) in CINs from Q175 mice (crossed with Thy1-ChR2 mice) are strongly reduced by ranolazine. The acquired dependence of distal cortical inputs on NaP currents in Q175 mice results from an upregulation of the NaP current, which was also evidenced by bAPs invading farther out into the CINs’ dendritic arbor (Tanimura et al., 2016). This elevated excitability in the Q175 mouse is probably a homeostatic response aimed at elevating the postsynaptic sensitivity to the remaining synaptic contents after the loss of afferent cortical and thalamic inputs observed in HD mouse models (Deng et al., 2013). Because thalamostriatal inputs are also altered in models of Parkinson’s disease (PD) (Aceves Buendia et al., 2019; Parker et al., 2016; Tanimura et al., 2019), future work should determine whether the excitability of CIN dendrites is altered in these models, as well. Importantly, dendrites are positioned at the interface between synaptic inputs and the intrinsic properties of CIN. While changes in synaptic transmission and intrinsic excitability of CINs have received attention in models of PD, HD and other movement disorders (Abudukeyoumu et al., 2019; Aceves Buendia et al., 2019; Choi et al., 2020; Ding et al., 2006, 2011; Eskow Jaunarajs et al., 2015; Mallet et al., 2019; Paz et al., 2021; Pisani et al., 2007; Plotkin and Goldberg, 2019; Poppi et al., 2021; Tanimura et al., 2019; Tubert and Murer, 2021), studying alterations in dendritic excitability in these models is a complementary approach that remains largely unchartered territory.

The membrane channels that are expressed in CINs (e.g., Nav, HCN, Kv4, etc.) are targets for neuromodulation under normal, healthy circumstances (Deng et al., 2007; Helseth et al., 2021; Maurice, 2004; Song and Surmeier, 1996). Therefore, it is likely that the dendritic excitability of CINs can be modulated under physiological conditions, as well, as a way to adjust the tuning of CINs to their afferent inputs. Conversely, if boosting and resonances are present primarily proximally, the selective tuning of CINs to subhertz slow-wave oscillations is probably stronger for thalamic inputs (that terminate proximally) than for cortical inputs. Future experiments can address this issue by testing how silencing the PfN affects the observed entrainment of TANs to slow-waves (Figure 6).

In summary, we have used a new optogenetics-based approach, complemented by other electrophysiological and imaging approaches, to demonstrate that the spatial localization of active dendritic conductances that endow CIN dendrites with quasi-linear filtering properties corresponds to the spatial distribution of their two main afferent excitatory inputs. This matching up of presynaptic terminals with post-synaptic excitability, probably contributes to the capacity of CINs to respond differentially to cortical and thalamic inputs. Sensorimotor information of cortical origin seems to be integrated continuously in a moment-by-moment fashion. In contrast, thalamic inputs lead to abrupt pauses in TAN firing, even in response to internally-generated brain-states (e.g., during slow-wave activity). The possibility that dendritic arbors adapt to the spatiotemporal patterns of afferent inputs is likely an important principle of neural computation that deserves further attention, particularly in the framework of autonomously active neurons such at CINs, SNr neurons and other basal ganglia pacemakers.

## Materials and Methods

| Key Resources Table |  |  |  |  |
| --- | --- | --- | --- | --- |
| Reagent type (species) or resource | Designation | Source or reference | Identifiers | Additional information |
| Genetic reagent (M. musculus) | B6;129S6-Chattm2(cre)Lowl/J | The Jackson Laboratory | Strain #:006410<br>RRID:IMSR_JAX:006410 | ChAt-ires-CRE |
| Genetic reagent (M. musculus) | B6;129S-Gt(ROSA)26Sortm32(CAG-COP4*H134R/EYFP)Hze/J | The Jackson Laboratory | Strain #:012569<br>RRID:IMSR_JAX:012569 | Ai32-conditional ChR2 |
| Genetic reagent (M. musculus) | STOCK Slc17a6tm2(cre)Lowl/J | The Jackson Laboratory | Strain #:016963<br>RRID:IMSR_JAX:016963 | Vglut2-ires-cre |
| Chemical compound, drug | Mecamylamine hydrochloride | Sigma-Aldrich | Lot # 019M4108V<br>CAS: 826-39-1 |  |
| Chemical compound, drug | Atropine sulfate salt monohydrate | Sigma-Aldrich | Lot # BCBH8339V<br>CAS No.: 5908-99-6 |  |
| Chemical compound, drug | SR 95531 hydrobromide (Gabazine) | Hello Bio | CAS: 104104-50-9 |  |
| Chemical compound, drug | DNQX | TOCRIS | CAS: 2379-57-9 |  |
| Chemical compound, drug | D-AP5 | Hello Bio | CAS:79055-68-8 |  |
| Chemical compound, drug | CGP 55845 hydrochloride | Hello Bio | CAS: 149184-22-5 |  |
| Chemical compound, drug | Dihydro-β-erythroidine hydrobromide | TOCRIS | CAS: 29734-68-7 | |
| Chemical compound, drug | Ranolazine dihydrochloride | Sigma-Aldrich | CAS Number: 95635-56-6 | Product Number R6152 |
| Chemical compound, drug | ZD7288 | MedChemExpress | CAS No. : 133059-99-1 | Synonyms: ICI D7288 |
| Chemical compound, drug | Tetrodotoxin citrate | Hello Bio | CAS: 18660-81-6 |  |
| Chemical compound, drug | 4-Aminopyridine | Sigma-Aldrich | CAS Number: 504-24-5 |  |
| Chemical compound, drug | Phosphate buffered saline tablets | Sigma-Aldrich | MDL number: MFCD00131855 | Product Number P4417 |
| Chemical compound, drug | Paraformaldehyde | Sigma-Aldrich | CAS Number: 30525-89-4 |  |
| Chemical compound, drug | XYLAZINE AS HYDROCHLORIDE | EUROVET ANIMAL HEALTH B.V | CAS: 082-91-92341-00 |  |
| Chemical compound, drug | CLORKETAM | VETOQUINOL | CAS: 1867-66-9 |  |
| Chemical compound, drug | Meloxicam | Chanelle Pharmaceuticals Manufacturing Ltd | CAS Number: 71125-38-7 |  |
| Chemical compound, drug | isoflurane | Primal Critical Care | CAS Number: 26675-46-7 |  |
| Software | MATLAB | MathWorks | RRID:SCR_001622 | Data analysis |
| Software | XPPAUT | X-Windows Phase Plane plus Auto | RRID:SCR_001996 | Modelling |
| Software | WinWCP | University of Strathclyde Glasgow | RRID:SCR_014713 | Data acquisition |
| Software | FEMTOsmart Software: MESc | FEMTONICS | RRID:SCR_018309 | 2P Data acquisition |
| Software | NIS-Elements Basic Research | Nikon instruments | RRID:SCR_002776 | Confocal images acquisition |
| Software | Signal 6 | Cambridge Electronic Design | RRID:SCR_017081 |  |
| Software | AlphaLab SnR | Alpha-Omega Engineering | https://www.alphaom-ega-eng.com/ |  |
| Software | Electrode Positioning System | Alpha-Omega Engineering | https://www.alphaom-ega-eng.com/ |  |
| Others | Model 940 Small Animal Stereotaxic Instrument with Digital Display Console | Kopf Instruments | https://kopfinstruments.com/product/model-940-small-animal-stereotaxic-instrument-with-digital-display-console/ |  |
| Others | Sound-attenuating room | IAC acoustics | https://www.iacacoustics.com/ |  |
| Others | Glass-coated Tungsten electrodes | Alpha-Omega Engineering | https://www.alphaomega-eng.com/ |  |
| Others | Pre-gelled surface electrodes | Rhythmlink International | https://rhythmlink.com/ |  |
| Others | subdermal needle electrodes | Rhythmlink International | https://rhythmlink.com/ |  |
| Others | Infra-red camera | Norpix Inc., Canada | https://www.norpix.com/ |  |

### Ethics statement

All experimental protocols were conducted in accordance with the National Institutes of Health Guide for the Care and Use of Laboratory Animals, and with the Hebrew University guidelines for the use and care of laboratory animals in research. The experiments adhered to, received prior written approval from and were supervised by the Institutional Animal Care and Use Committee of the Faculty of Medicine, under protocols: MD-16-13518-4 (H.B.) and MD-18-15657-3 (J.A.G.).

### Mice

Homozygous Ai32 (RCL-ChR2(H134R)/EYFP) mice (The Jackson laboratory [Jax] stock: 012569) that express floxed ChR2 and an EYFP fusion protein under the CAG promoter were crossed with homozygous ChAT-IRES-Cre (*Δ*neo) mice the express Cre recombinase under the choline acetyltransferase promoter (Jax stock: 031661). The ChAT-ChR2 offspring (4-8 weeks old/both sexes) were used for the majority of experiments. To investigate corticostriatal transmission, we used homozygous transgenic Thy1-ChR2 mice (B6.Cg-Tg (Thy1-COP4/EYFP) 18Gfng/1, Jax stock: 007612), that express ChR2 under the Thy1 promoter in cortical neurons (Arenkiel et al., 2007). To investigate thalamostriatal transmission, we used Vglut2-ires-Cre mice (Jax stock: 016963).

### Non-Human Primates (NHPs)

Data were obtained from two healthy, young adult, female vervet monkeys. The monkeys were habituated to sleeping in a primate chair, positioned in a dark, double-walled sound-attenuating room. The primate chair restrained the monkeys’ hand and body movements but otherwise allowed them to be in a position similar to their natural (sitting) sleeping posture. Detailed sleep habituation, surgery and sleep staging were reported previously (Mizrahi-Kliger et al., 2018). For extracellular recordings, the monkeys’ heads were immobilized with a head holder and eight glass-coated tungsten microelectrodes were advanced the dorsal striatum. Electrical signals were amplified with a gain of 20, filtered using a 0.075 Hz (2 pole) to 10 kHz (3 pole) Butterworth filter and sampled at 44 kHz by a 16-bit analog/digital converter. Spiking activity was sorted online using a template matching algorithm. The striatum was identified based on its stereotaxic coordinates according to MRI imaging and primate atlas data (Martin and Bowden, 2000). Tonically active neurons (TANs) and spiny projection neurons (SPNs) were identified using real-time assessment of their electrophysiological features. Spiking and local field potentials (LFP) were recorded only for identified recording sites with stable recording quality (i.e., where single-neuron spiking yielded an average isolation score ≥0.85).

We followed established procedures for slow wave detection in the LFP (Riedner et al., 2007). Briefly, the LFP signal was filtered at the 0.5-4 Hz range and putative slow wave events whose duration was 0.25-2 seconds were kept for further analysis. Next, the slow wave peaks were sorted according to their amplitude. Artifacts were removed by discarding all events whose amplitude exceeded 5 standard deviations above the mean. Finally, conditional firing rate analysis was only performed on 30% of the slow wave events with the highest amplitude. Conventional procedures were also used for sleep spindle detection in the LFP (Sela et al., 2016, p. 2016). Briefly, the detection algorithm was only used for striatal sites whose LFP showed significant 10-17 Hz (spindle range) activity. LFP was filtered at the 10-17 Hz range and the Hilbert transform was then used to extract the instantaneous amplitude. Events exceeding 3 standard deviations above the mean were deemed potential spindle events, and a threshold of one half of a standard deviation above the mean was used to detect the start and end point of an individual sleep spindle. A potential sleep spindle was defined as such only if it lasted 0.5-3 seconds, and provided it did not exhibit a relatively high (more than 4.5 standard deviations above the mean) amplitude in a control 20-30 Hz range. The spindle data were obtained using a 4-pole 4-25 Hz Butterworth filter.

Histology: An 8 weeks old male Thy1-ChR2 mouse was deeply anesthetized with a terminal dose of ketamine-xylazine followed by perfusion through the heart of cold PBS and 4% PFA. The removed brain was kept overnight at 4°C in 4% PFA. The next day, the brain was washed 3 × 15 minutes before 50 µm coronal slices of the PfN were cut with a vibratome (Leica VT1000S). VECTASHIELD (Vector Laboratories) was applied onto mounted slices to protect from bleaching. Coronal slices of the PfN were imaged using confocal microscope (Nikon A1R) using 10x lens and a 20x oil immersion lens to visualize constitutive EYFP expression.

### Stereotaxic viral inoculation in Vglut2-ires-Cre mice

Mice were deeply anesthetized with isoflurane in a non-rebreathing system (2.5% induction, 1-1.5% maintenance) and placed in a stereotaxic frame (Kopf Instruments, Tujunga, CA). Temperature was maintained at 35°C with a heating pad, artificial tears were applied to prevent corneal drying, and animals were hydrated with a bolus of injectable saline (10 ml/kg) mixed with analgesic (2.5 mg/kg Meloxicam). Stereotaxic injections into caudal intralaminar nuclei of thalamus were performed under aseptic conditions. Adeno-associated viruses (AAV) serotype 9 carrying double-floxed fusion genes for hChR2 (E123A) and EYFP under an EF1a promoter (University of Pennsylvania Vector Core, Addgene #35507) were used to transfect PfN neurons. Injection coordinates were from Bregma: lateral, 0.65 mm; posterior, 2.3 mm; and 3.35 mm depth from surface of brain (Rehani et al., 2019). A small hole was bored into the skull with a micro drill bit and a glass pipette was slowly inserted at the PfN coordinates. To minimize backflow, solution was slowly injected; a total volume of 250 nl (>2.5×1012 GC/ml) of the AAV constructs was injected over a period of approximately 1.5 min and the pipette was left in place for 5 min before slowly retracting it. Slice physiology experiments were conducted 2-3 weeks after surgery.

### Slice preparation

Mice were deeply anesthetized with ketamine (200 mg/kg)–xylazine (23.32 mg/kg) and perfused transcardially with ice-cold-modified artificial cerebrospinal fluid (ACSF) bubbled with 95% O_2_–5% CO_2_, and containing (in mM) 2.5 KCl, 26 NaHCO_3_, 1.25 Na2HPO4, 0.5 CaCl_2_, 10 MgSO_4_, 0.4 ascorbic acid, 10 glucose and 210 sucrose. The brain was removed, and 275 µm thick sagittal slices containing the striatum were cut in ice-cold-modified ACSF. Slices were then submerged in ACSF, bubbled with 95% O_2_–5% CO_2_, containing (in mM) 2.5 KCl, 126 NaCl, 26 NaHCO_3_, 1.25 Na_2_HPO_4_, 2 CaCl_2_, 2 MgSO_4_ and 10 glucose, and stored at room temperature for at least 1 h prior to recording.

### Electrophysiological recording

The slices were transferred to the recording chamber mounted on a Zeiss Axioskop 60X, 0.9 NA fixed-stage microscope and perfused with oxygenated ACSF at 31ºC. During the optogenetic stimulation experiments, in order to guarantee that the effects we measured were generated post-synaptically, the ACSF solution contained (in µM) 10 DNQX to block AMPA receptors, 50 D-APV to block NMDA receptors, 10 gabazine (SR95531) to block GABA_A_ receptors, - 2 CGP55845 to block GABA_B_ receptors, 10 atropine to block muscarinic ACh receptors, and 10 mecamylamine to block nicotinic ACh receptors. In the experiments in which optogenetics were used to stimulate cortical or thalamic input we used the same blockers, except for DNQX which was left out. An Olympus 40X, 0.8 NA water-immersion objective with a 26.5 mm field number (FN) was used to examine the slice using standard infrared differential interference contrast video microscopy. Patch pipette resistance was typically 4-5 MΩ when filled with recording solutions. The junction potential estimated at 7-8 mV was not corrected. In EPSC measurements, the intracellular solution contained (in mM): 127.5 CsCH_3_SO_3_, 7.5 CsCl, 10 HEPES, 10 TEA-Cl, 4 phosphocreatine disodium, 0.2 EGTA, 0.21 Na_2_GTP, and 2 Mg_1.5_ATP (pH=7.3 with CsOH, 280-290 mOsm/kg). In the Ca^2+^ imaging experiments (see below) the internal solution contained (in mM) 135 K-gluconate, 5 KCl, 2.5 NaCl, 5 Na-phosphocreatine, 10 HEPES, 0.1 fluo-4 (Molecular Probes), 0.1 Alexa Fluor 568 (for morphological visualization, Molecular Probes), 0.21 Na_2_GTP, and 2 Mg_1.5_ATP, pH 7.3 with KOH (280–290 mOsm/kg). In all other experiments, the intracellular solution contained (in mM) 135.5 KCH_3_SO_3_, 5 KCl, 2.5 NaCl, 5 Na-phosphocreatine, 10 HEPES, 0.2 EGTA, 0.21 Na_2_GTP, and 2 Mg_1.5_ATP, pH 7.3 with KOH (280–290 mOsm/kg). Electrophysiological recordings were obtained with a MultiClamp 700B amplifier (Molecular Devices, Sunnyvale, CA). Signals were filtered at 10 kHz online, digitized at 10 or 20 kHz and logged onto a personal computer with the Signal 6 software (Cambridge Electronic Design, Cambridge, UK). All reagents, drugs and toxins were purchased from either Merck/Sigma-Aldrich (Darmstadt, Germany), Tocris Biosciences (Bristol, UK), MedChemExpress (Monmouth Junction, NJ, USA) or HelloBio (Bristol, UK).

### Voltage-perturbation experiments

CINs were held at either –55 mV or –70 mV and were given an 83 second-long voltage command structured as a concatenated sequence of sinusoids from a discrete set of frequencies ranging from 0.2 to 20 Hz with an amplitude of 2 mV (3 or 5 seconds per frequency, such that each frequency was represented by an integer multiple of its fundamental period). Phase shifts between the voltage sinusoidal and the somatic current response were determined by the location of the peak in the cross-correlation function (CCF) of the two traces (whose units are mV·pA), for each stimulation frequency and for each illumination condition. The impedance at each frequency, |*Z*(*f*)|, was calculated from the maximal amplitude of the CCF as |*Z*(*f*)| = (2 mV)^2^/max(CCF) (so that its units are GΩ).

### Optogenetic stimulation

Optogenetic stimulation was performed with blue-light (470 nm) LED illumination via the objective (Mightex, Toronto, ON, Canada). We used two spatial illumination regimes: a) *proximal illumination* wherein an opaque disk with a central pinhole was placed in the back focal plane of the 60X water-immersion objective such that a ∼130 μm diameter region around the soma was illuminated (Tiroshi and Goldberg, 2019), thereby targeting the soma and proximal dendrites; and b) *full-field illumination* of the entire slice with a 5X air objective which excites the soma and the entire dendritic field. In all experiments, LED light intensity was chosen such that stimulation generated comparable current responses for both regimes. We used the same sequence of sinusoids described above, only this time the voltage driving the LED was modulated (the minimal voltage was the LED’s voltage threshold, 40 mV). The phase delays were again calculated according to the latency of the peak of the CCF between the LED voltage command and somatic current.

Note, that the phases were corrected by 0.5 (i.e., by π in radians) due to the fact that the ChR2 inward current is in antiphase with the LED’s voltage command. The amplitude response was calculated from the peak value of the CCF, normalized by the amplitude of the 470 nm LED command (i.e., 1V for proximal illumination and 0.1 V for the full-field illumination, so that its units are picoamperes).

To activate the excitatory synaptic inputs in the Thy1-ChR2 and in the Vglut2-mice a full-field 470 nm LED 1 ms-long pulses were used with GABA, ACh and NMDA receptor blockers in the ACSF. For EPSPs we average 25 trials (3 s intervals, and trials with spikes were omitted). Paired pulse ratio (PPR) measurements consisted of 64 trials of two pulses (100 ms apart, 3 s interval). The mean EPSC amplitude was calculated as the difference between the mean peak current and the mean baseline current that preceded the pulse. PPR was the ratio of the second mean EPSC to the first mean EPSC. To demonstrate that the EPSCs were monosynaptic they were recorded before and after application of 1 µM TTX and 100 µM 4-AP (Petreanu et al., 2009).

To estimate the kinetics of the ChR2 currents, brief 1ms-long 470 nM LED pulses (1V for proximal and 0.1 V for the full-field illumination) were repeated 250 times and the resulting average current response was measured, and fit with an alpha function

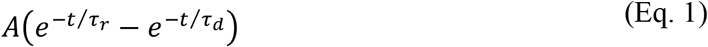

to estimate the *τ* _*r*_ and *τ*_*d*_, the rise and decay times, respectively.

### Two-photon laser scanning microscopy (2PLSM)

The two-photon excitation source was a Chameleon Vision II tunable Ti:Sapphire pulsed laser (Coherent, Santa Clara, CA, USA) tuned to 820 nm. The images were collected with the Femto2D system (Femtonics, Budapest, Hungary) which includes two 3 mm galvo-scanners, one gated GaAsP and one multi-alkaline non-descanned photomultiplier tube for imaging fluo-4 and Alexa Fluor, respectively. Z-stacks of optical sections (spaced 2 µm apart) were collected using 0.2 µm pixels and 15 µs dwell times. Optical and electrophysiological data were obtained using the software package MES (Femtonics), which also integrates the control of all hardware units in the microscope. The software automates and synchronizes the imaging signals and electrophysiological protocols. Data was extracted from the MES package to personal computers using custom-made code in MATLAB. We recorded spontaneously occurring bAPs with line scans at various distances measured radially from the tip of the soma. Spike triggered averages of the calcium measurements (*ΔF/F*_*0*_) were estimated and an alpha-function (Eq. 1) was fit to them. The value of the peak of the fitted alpha-function was used as a measure of the size of the spontaneous bAP at that location.

### Drugs and Reagents

TTX was used to block voltage-activated Na^+^ currents. Ranolazine was used to block NaP. ZD7288 was used to block the HCN current. Gabazine (SR-95531) and CGP 55845 were used to block GABA_A_ and GABA_B_ receptors, respectively. 4-Aminopyridine (4-AP) was used to enable optogenetically-driven monosynaptic release in the presence of TTX. All reagents, drugs and toxins were purchased from either Merck/Sigma-Aldrich (Darmstadt, Germany), Tocris Biosciences (Bristol, UK) or HelloBio (Bristol, UK).

### Data analysis and statistics

Data were analyzed and curve fitting was performed using custom code in Matlab (MathWorks, Natick, MA, USA). Simulations of the nonlinear dendrite (Appendix 2) was were performed using XPPAUT (Ermentrout, 2002). The nonparametric two-tailed Wilcoxon signed-rank test was used for matched samples and the Wilcoxon rank-sum test was used for independent samples. The parametric ANCOVA test was used to test significant changes in the amplitude and phase curves as a function of the natural logarithm of the frequencies (an operation which spreads out this independent parameter more uniformly). Null hypotheses were rejected if the P-value was below 0.05.

For the TAN and SPN locking to slow wave or spindle peak analysis, confidence intervals (at P-value of 0.01) were calculated based on the distribution of conditional firing rates 1 s before and after the slow wave peak.

### Parameter fitting to the phase delay

In our previous study (Tiroshi and Goldberg, 2019), we modeled the dendritic arbor as a semi-infinite cable with a homogeneous quasi-linear membrane (i.e., the current density of each nonlinearity is constant along the dendrite). When a segment of length *r* (measured in units of the dendrite’s space constant) from the soma is activated with a sinusoidal current injection, the dendritic phase delay is given by

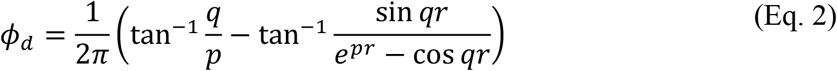

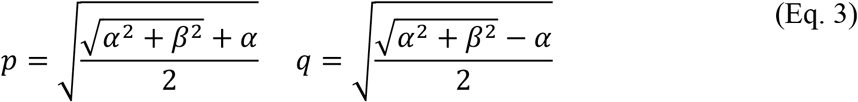

*α* and β are functions of frequency and are determined by the linearization of the dendritic nonlinearities as explained in Goldberg et al. (2007) and Remme and Rinzel (2011), with a negative amplifying parameter, *µ*_*n*_, and a positive resonance parameter, *µ*_*h*_ (See Appendix 1).

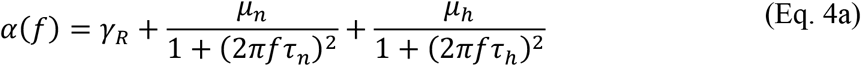

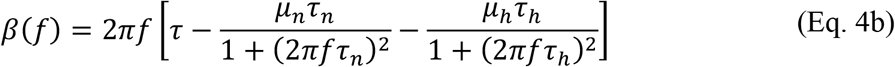

Additional parameters include total dendritic conductance (relative to leak), γ,, the membrane time constant τ, and the time constants representing the kinetics of the nonlinear dendritic conductances, as explained in Goldberg et al. (2007) and in Remme and Rinzel (2011). In some cases we only used the amplifying parameter in the fit (e.g., Figure 1C), and in Figure 3C, we used a Eq. 2 in the case of a passive dendrite for which α*(f)* = 1 and β*(f) =*2π*f*τ.

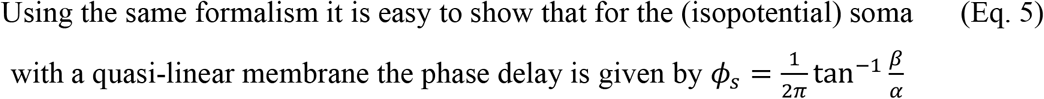

corresponding to an amplitude that is equal to (α^2^ + β^2^)^-1/2^ up to a scaling factor. In the main text, we point out that when *µ*_*n*_ becomes less negative *ϕ*_s_ is reduced. This is because when *µ*_*n*_ becomes less negative, *α* is increased (Eq. 4a) and *β* is decreased (Eq. 4b).

The amplitude attenuation and phase delays generated by the ChR2 kinetics are calculated from the Fourier transform of the alpha function (Eq. 1) and are given by

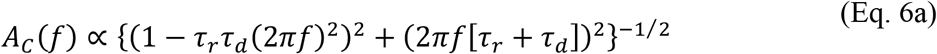

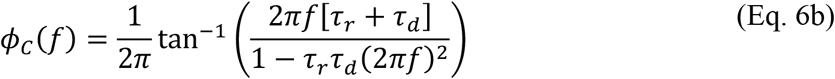

## Acknowledgments

This work was funded by a European Research Council (ERC) Consolidator Grant (no. 646886) to J.A.G. and grants from the Israel Science Foundation (no. 2051/20), German Research Foundation (CRC TRR295), Israel-China Binational Science Foundation and the Silverstein Foundation to H.B. We would like to thank Ronit Cherki, Tamar Licht and Anatoly Shapochnikov for excellent technical assistance.

## References

Abudukeyoumu N, Hernandez-Flores T, Garcia-Munoz M, Arbuthnott GW. 2019. Cholinergic modulation of striatal microcircuits. Eur J Neurosci 49:604–622. doi:10.1111/ejn.13949

Aceves Buendia J de J, Tiroshi L, Chiu W, Goldberg JA. 2019. Selective remodeling of glutamatergic transmission to striatal cholinergic interneurons after dopamine depletion. Eur J Neurosci 49:824–833. doi:10.1111/ejn.13715

Alzheimer C, Schwindt P, Crill W. 1993. Modal gating of Na+ channels as a mechanism of persistent Na+ current in pyramidal neurons from rat and cat sensorimotor cortex. J Neurosci 13:660–673. doi:10.1523/JNEUROSCI.13-02-00660.1993

Aoki S, Liu AW, Akamine Y, Zucca A, Zucca S, Wickens JR. 2018. Cholinergic interneurons in the rat striatum modulate substitution of habits. Eur J Neurosci 47:1194–1205. doi:10.1111/ejn.13820

Aosaki T, Kimura M, Graybiel AM. 1995. Temporal and spatial characteristics of tonically active neurons of the primate’s striatum. J Neurophysiol 73:1234–1252.

Aosaki T, Tsubokawa H, Ishida A, Watanabe K, Graybiel AM, Kimura M. 1994. Responses of tonically active neurons in the primate’s striatum undergo systematic changes during behavioral sensorimotor conditioning. J Neurosci 14:3969–3984.

Apicella P, Scarnati E, Schultz W. 1991. Tonically discharging neurons of monkey striatum respond to preparatory and rewarding stimuli. Exp Brain Res 84:672–675.

Arenkiel BR, Peca J, Davison IG, Feliciano C, Deisseroth K, Augustine GJ, Ehlers MD, Feng G. 2007. In vivo light-induced activation of neural circuitry in transgenic mice expressing channelrhodopsin-2. Neuron 54:205–218. doi:10.1016/j.neuron.2007.03.005

Assous M. 2021. Striatal cholinergic transmission. Focus on nicotinic receptors’ influence in striatal circuits. Eur J Neurosci 53:2421–2442. doi:10.1111/ejn.15135

Assous M, Dautan D, Tepper JM, Mena-Segovia J. 2019. Pedunculopontine Glutamatergic Neurons Provide a Novel Source of Feedforward Inhibition in the Striatum by Selectively Targeting Interneurons. J Neurosci 39:4727–4737. doi:10.1523/JNEUROSCI.2913-18.2019

Assous M, Kaminer J, Shah F, Garg A, Koós T, Tepper JM. 2017. Differential processing of thalamic information via distinct striatal interneuron circuits. Nature Communications 8:15860. doi:10.1038/ncomms15860

Beatty Joseph A., Song SC, Wilson CJ. 2015. Cell-type-specific resonances shape the responses of striatal neurons to synaptic input. Journal of Neurophysiology 113:688–700. doi:10.1152/jn.00827.2014

Bennett BD, Callaway JC, Wilson CJ. 2000. Intrinsic membrane properties underlying spontaneous tonic firing in neostriatal cholinergic interneurons. J Neurosci 20:8493–8503. doi:20/22/8493 [pii]

Bennett BD, Wilson CJ. 1999. Spontaneous activity of neostriatal cholinergic interneurons in vitro. J Neurosci 19:5586–5596.

Berger T, Larkum ME, Lüscher H-R. 2001. High I h Channel Density in the Distal Apical Dendrite of Layer V Pyramidal Cells Increases Bidirectional Attenuation of EPSPs. Journal of Neurophysiology 85:855–868. doi:10.1152/jn.2001.85.2.855

Bradfield LA, Bertran-Gonzalez J, Chieng B, Balleine BW. 2013. The Thalamostriatal Pathway and Cholinergic Control of Goal-Directed Action: Interlacing New with Existing Learning in the Striatum. Neuron 79:153–166. doi:10.1016/j.neuron.2013.04.039

Brown RE, Basheer R, McKenna JT, Strecker RE, McCarley RW. 2012. Control of Sleep and Wakefulness. Physiological Reviews 92:1087–1187. doi:10.1152/physrev.00032.2011

Buzsáki G, Anastassiou CA, Koch C. 2012. The origin of extracellular fields and currents — EEG, ECoG, LFP and spikes. Nat Rev Neurosci 13:407–420. doi:10.1038/nrn3241

Carter AG, Sabatini BL. 2004. State-Dependent Calcium Signaling in Dendritic Spines of Striatal Medium Spiny Neurons. Neuron 44:483–493. doi:10.1016/j.neuron.2004.10.013

Choi SJ, Ma TC, Ding Y, Cheung T, Joshi N, Sulzer D, Mosharov EV, Kang UJ. 2020. Alterations in the intrinsic properties of striatal cholinergic interneurons after dopamine lesion and chronic L-DOPA. eLife 9:e56920. doi:10.7554/eLife.56920

Day M, Wokosin D, Plotkin JL, Tian X, Surmeier DJ. 2008. Differential Excitability and Modulation of Striatal Medium Spiny Neuron Dendrites. Journal of Neuroscience 28:11603–11614. doi:10.1523/JNEUROSCI.1840-08.2008

Deng P, Zhang Y, Xu ZC. 2007. Involvement of I(h) in dopamine modulation of tonic firing in striatal cholinergic interneurons. J Neurosci 27:3148–3156. doi:27/12/3148 [pii] 10.1523/JNEUROSCI.5535-06.2007

Deng YP, Wong T, Bricker-Anthony C, Deng B, Reiner A. 2013. Loss of corticostriatal and thalamostriatal synaptic terminals precedes striatal projection neuron pathology in heterozygous Q140 Huntington’s disease mice. Neurobiol Dis 60:89–107. doi:10.1016/j.nbd.2013.08.009

Ding J, Guzman JN, Tkatch T, Chen S, Goldberg JA, Ebert PJ, Levitt P, Wilson CJ, Hamm HE, Surmeier DJ. 2006. RGS4-dependent attenuation of M4 autoreceptor function in striatal cholinergic interneurons following dopamine depletion. Nat Neurosci 9:832–842. doi:10.1038/nn1700

Ding JB, Guzman JN, Peterson JD, Goldberg JA, Surmeier DJ. 2010. Thalamic Gating of Corticostriatal Signaling by Cholinergic Interneurons. Neuron 67:294–307. doi:10.1016/j.neuron.2010.06.017

Ding Y, Won L, Britt JP, Lim SAO, McGehee DS, Kang UJ. 2011. Enhanced striatal cholinergic neuronal activity mediates L-DOPA-induced dyskinesia in parkinsonian mice. Proceedings of the National Academy of Sciences 108:840–845. doi:10.1073/pnas.1006511108

Doig NM, Magill PJ, Apicella P, Bolam JP, Sharott A. 2014. Cortical and Thalamic Excitation Mediate the Multiphasic Responses of Striatal Cholinergic Interneurons to Motivationally Salient Stimuli. Journal of Neuroscience 34:3101–3117. doi:10.1523/JNEUROSCI.4627-13.2014

Ermentrout B. 2002. Simulating, Analyzing, and Animating Dynamical Systems: A Guide to XPPAUT for Researchers and Students. Philadelphia: Society for Industrial and Applied Mathematics.

Eskow Jaunarajs KL, Bonsi P, Chesselet MF, Standaert DG, Pisani A. 2015. Striatal cholinergic dysfunction as a unifying theme in the pathophysiology of dystonia. Progress in Neurobiology 127–128:91–107. doi:10.1016/j.pneurobio.2015.02.002

Goldberg JA, Deister CA, Wilson CJ. 2007. Response Properties and Synchronization of Rhythmically Firing Dendritic Neurons. Journal of Neurophysiology 97:208–219. doi:10.1152/jn.00810.2006

Goldberg JA, Ding JB, Surmeier DJ. 2012. Muscarinic Modulation of Striatal Function and Circuitry In: Fryer AD, Christopoulos A, Nathanson NM, editors. Muscarinic Receptors, Handbook of Experimental Pharmacology. Berlin, Heidelberg: Springer Berlin Heidelberg. pp. 223–241. doi:10.1007/978-3-642-23274-9_10

Goldberg JA, Kats SS, Jaeger D. 2003. Globus Pallidus Discharge Is Coincident with Striatal Activity during Global Slow Wave Activity in the Rat. J Neurosci 23:10058–10063. doi:10.1523/JNEUROSCI.23-31-10058.2003

Goldberg JA, Reynolds JN. 2011. Spontaneous firing and evoked pauses in the tonically active cholinergic interneurons of the striatum. Neuroscience 198:27–43. doi:10.1016/j.neuroscience.2011.08.067

Goldberg JA, Teagarden MA, Foehring RC, Wilson CJ. 2009. Nonequilibrium Calcium Dynamics Regulate the Autonomous Firing Pattern of Rat Striatal Cholinergic Interneurons. Journal of Neuroscience 29:8396–8407. doi:10.1523/JNEUROSCI.5582-08.2009

Goldberg JA, Wilson CJ. 2010. The Cholinergic Interneurons of the Striatum: Intrinsic Properties Underlie Multiple Discharge Patterns. In: Steiner H, Tseng KY, editors. Handbook of Basal Ganglia Structure and Function. London: Academic Press. pp. 133–149.

Goldberg JA, Wilson CJ. 2005. Control of Spontaneous Firing Patterns by the Selective Coupling of Calcium Currents to Calcium-Activated Potassium Currents in Striatal Cholinergic Interneurons. Journal of Neuroscience 25:10230–10238. doi:10.1523/JNEUROSCI.2734-05.2005

Gradinaru V, Mogri M, Thompson KR, Henderson JM, Deisseroth K. 2009. Optical deconstruction of parkinsonian neural circuitry. Science 324:354–359. doi:10.1126/science.1167093

Graybiel A, Aosaki T, Flaherty A, Kimura M. 1994. The basal ganglia and adaptive motor control. Science 265:1826–1831. doi:10.1126/science.8091209

Gritton HJ, Howe WM, Romano MF, DiFeliceantonio AG, Kramer MA, Saligrama V, Bucklin ME, Zemel D, Han X. 2019. Unique contributions of parvalbumin and cholinergic interneurons in organizing striatal networks during movement. Nat Neurosci 22:586–597. doi:10.1038/s41593-019-0341-3

Harnett MT, Magee JC, Williams SR. 2015. Distribution and Function of HCN Channels in the Apical Dendritic Tuft of Neocortical Pyramidal Neurons. Journal of Neuroscience 35:1024–1037. doi:10.1523/JNEUROSCI.2813-14.2015

Häusser M. 2021. Optogenetics — The Might of Light. N Engl J Med 385:1623–1626. doi:10.1056/NEJMcibr2111915

Helseth AR, Hernandez-Martinez R, Hall VL, Oliver ML, Turner BD, Caffall ZF, Rittiner JE, Shipman MK, King CS, Gradinaru V, Gerfen C, Costa-Mattioli M, Calakos N. 2021. Cholinergic neurons constitutively engage the ISR for dopamine modulation and skill learning in mice. Science 372:eabe1931. doi:10.1126/science.abe1931

Higgs MH, Wilson CJ. 2017. Measurement of phase resetting curves using optogenetic barrage stimuli. Journal of Neuroscience Methods 289:23–30. doi:10.1016/j.jneumeth.2017.06.018

Howe M, Ridouh I, Allegra Mascaro AL, Larios A, Azcorra M, Dombeck DA. 2019. Coordination of rapid cholinergic and dopaminergic signaling in striatum during spontaneous movement. eLife 8:e44903. doi:10.7554/eLife.44903

Hutcheon B, Miura RM, Puil E. 1996. Subthreshold membrane resonance in neocortical neurons. Journal of Neurophysiology 76:683–697. doi:10.1152/jn.1996.76.2.683

Hutcheon B, Yarom Y. 2000. Resonance, oscillation and the intrinsic frequency preferences of neurons. Trends in Neurosciences 23:216–222. doi:10.1016/S0166-2236(00)01547-2

Johansson Y, Silberberg G. 2020. The Functional Organization of Cortical and Thalamic Inputs onto Five Types of Striatal Neurons Is Determined by Source and Target Cell Identities. Cell Reports 30:1178–1194.e3. doi:10.1016/j.celrep.2019.12.095

Joshua M, Adler A, Mitelman R, Vaadia E, Bergman H. 2008. Midbrain Dopaminergic Neurons and Striatal Cholinergic Interneurons Encode the Difference between Reward and Aversive Events at Different Epochs of Probabilistic Classical Conditioning Trials. Journal of Neuroscience 28:11673–11684. doi:10.1523/JNEUROSCI.3839-08.2008

Kawaguchi Y. 1993. Physiological, morphological, and histochemical characterization of three classes of interneurons in rat neostriatum. J Neurosci 13:4908–4923. doi:10.1523/JNEUROSCI.13-11-04908.1993

Kerr JND. 2004. Action Potential Timing Determines Dendritic Calcium during Striatal Up-States. Journal of Neuroscience 24:877–885. doi:10.1523/JNEUROSCI.4475-03.2004

Kim CK, Adhikari A, Deisseroth K. 2017. Integration of optogenetics with complementary methodologies in systems neuroscience. Nat Rev Neurosci 18:222–235. doi:10.1038/nrn.2017.15

Kimura M, Rajkowski J, Evarts E. 1984. Tonically discharging putamen neurons exhibit set-dependent responses. Proceedings of the National Academy of Sciences 81:4998–5001. doi:10.1073/pnas.81.15.4998

Koch C. 1984. Cable theory in neurons with active, linearized membranes. Biol Cybern 50:15–33. doi:10.1007/BF00317936

Kole MHP, Hallermann S, Stuart GJ. 2006. Single I h Channels in Pyramidal Neuron Dendrites: Properties, Distribution, and Impact on Action Potential Output. J Neurosci 26:1677–1687. doi:10.1523/JNEUROSCI.3664-05.2006

Kosillo P, Zhang Y-F, Threlfell S, Cragg SJ. 2016. Cortical Control of Striatal Dopamine Transmission via Striatal Cholinergic Interneurons. Cereb Cortex 26:4160–4169. doi:10.1093/cercor/bhw252

Lapper SR, Bolam JP. 1992. Input From the Frontal Cortex and the Nucleus To Cholinergic Interneurons in the Dorsal of the Rat 51:533–545.

Liu D, Dan Y. 2019. A Motor Theory of Sleep-Wake Control: Arousal-Action Circuit. Annu Rev Neurosci 42:27–46. doi:10.1146/annurev-neuro-080317-061813

Mahon S. 2001. Relationship between EEG Potentials and Intracellular Activity of Striatal and Cortico-striatal Neurons: an In Vivo Study under Different Anesthetics. Cerebral Cortex 11:360–373. doi:10.1093/cercor/11.4.360

Mallet N, Leblois A, Maurice N, Beurrier C. 2019. Striatal Cholinergic Interneurons: How to Elucidate Their Function in Health and Disease. Front Pharmacol 10:1488. doi:10.3389/fphar.2019.01488

Mamaligas AA, Barcomb K, Ford CP. 2019. Cholinergic Transmission at Muscarinic Synapses in the Striatum Is Driven Equally by Cortical and Thalamic Inputs. Cell Reports 28:1003–1014.e3. doi:10.1016/j.celrep.2019.06.077

Martin RF, Bowden DM. 2000. Primate brain maps: structure of the macaque brain. Elsevier.

Matityahu L, Malgady JM, Schirelman M, Johansson Y, Wilking JA, Silberberg G, Goldberg JA, Plotkin JL. 2022. A tonic nicotinic brake controls spike timing in striatal spiny projection neurons. eLife 11:e75829. doi:10.7554/eLife.75829

Matsumoto N, Minamimoto T, Graybiel AM, Kimura M. 2001. Neurons in the Thalamic CM-Pf Complex Supply Striatal Neurons With Information About Behaviorally Significant Sensory Events. Journal of Neurophysiology 85:960–976. doi:10.1152/jn.2001.85.2.960

Maurice N. 2004. D2 Dopamine Receptor-Mediated Modulation of Voltage-Dependent Na+ Channels Reduces Autonomous Activity in Striatal Cholinergic Interneurons. Journal of Neuroscience 24:10289–10301. doi:10.1523/JNEUROSCI.2155-04.2004

Mauro A, Conti F, Dodge F, Schor R. 1970. Subthreshold Behavior and Phenomenological Impedance of the Squid Giant Axon. Journal of General Physiology 55:497–523. doi:10.1085/jgp.55.4.497

McGuirt AF, Post MR, Pigulevskiy I, Sulzer D, Lieberman OJ. 2021. Coordinated Postnatal Maturation of Striatal Cholinergic Interneurons and Dopamine Release Dynamics in Mice. J Neurosci 41:3597–3609. doi:10.1523/JNEUROSCI.0755-20.2021

Mizrahi-Kliger AD, Kaplan A, Israel Z, Bergman H. 2018. Desynchronization of slow oscillations in the basal ganglia during natural sleep. Proc Natl Acad Sci USA 115:E4274–E4283. doi:10.1073/pnas.1720795115

Morris G, Arkadir D, Nevet A, Vaadia E, Bergman H. 2004. Coincident but distinct messages of midbrain dopamine and striatal tonically active neurons. Neuron 43:133–143. doi:10.1016/j.neuron.2004.06.012 S0896627304003551 [pii]

Nagel G, Szellas T, Huhn W, Kateriya S, Adeishvili N, Berthold P, Ollig D, Hegemann P, Bamberg E. 2003. Channelrhodopsin-2, a directly light-gated cation-selective membrane channel. Proc Natl Acad Sci USA 100:13940–13945. doi:10.1073/pnas.1936192100

Oswald MJ, Oorschot DE, Schulz JM, Lipski J, Reynolds JN. 2009. IH current generates the afterhyperpolarisation following activation of subthreshold cortical synaptic inputs to striatal cholinergic interneurons. J Physiol 587:5879–5897. doi:jphysiol.2009.177600 [pii] 10.1113/jphysiol.2009.177600

Parker PRL, Lalive AL, Kreitzer AC. 2016. Pathway-Specific Remodeling of Thalamostriatal Synapses in Parkinsonian Mice. Neuron 89:734–740. doi:10.1016/j.neuron.2015.12.038

Paz RM, Tubert C, Stahl AM, Amarillo Y, Rela L, Murer MG. 2021. Levodopa Causes Striatal Cholinergic Interneuron Burst-Pause Activity in Parkinsonian Mice. Mov Disord 36:1578–1591. doi:10.1002/mds.28516

Petreanu L, Mao T, Sternson SM, Svoboda K. 2009. The subcellular organization of neocortical excitatory connections. Nature 457:1142–5. doi:10.1038/nature07709

Pisani A, Bernardi G, Ding J, Surmeier DJ. 2007. Re-emergence of striatal cholinergic interneurons in movement disorders. Trends in Neurosciences 30:545–553. doi:10.1016/j.tins.2007.07.008

Plotkin JL, Day M, Surmeier DJ. 2011. Synaptically driven state transitions in distal dendrites of striatal spiny neurons. Nat Neurosci 14:881–888. doi:10.1038/nn.2848

Plotkin JL, Goldberg JA. 2019. Thinking Outside the Box (and Arrow): Current Themes in Striatal Dysfunction in Movement Disorders. Neuroscientist 25:359–379. doi:10.1177/1073858418807887

Poppi LA, Ho-Nguyen KT, Shi A, Daut CT, Tischfield MA. 2021. Recurrent Implication of Striatal Cholinergic Interneurons in a Range of Neurodevelopmental, Neurodegenerative, and Neuropsychiatric Disorders. Cells 10:907. doi:10.3390/cells10040907

Raz A, Feingold A, Zelanskaya V, Vaadia E, Bergman H. 1996. Neuronal synchronization of tonically active neurons in the striatum of normal and parkinsonian primates. J Neurophysiol 76:2083–2088.

Rehani R, Atamna Y, Tiroshi L, Chiu W-H,de Jesús Aceves Buendía J, Martins GJ, Jacobson GA, Goldberg JA. 2019. Activity Patterns in the Neuropil of Striatal Cholinergic Interneurons in Freely Moving Mice Represent Their Collective Spiking Dynamics. eNeuro 6:ENEURO.0351-18.2018. doi:10.1523/ENEURO.0351-18.2018

Reig R, Silberberg G. 2014. Multisensory Integration in the Mouse Striatum. Neuron 83:1200–1212. doi:10.1016/j.neuron.2014.07.033

Remme MWH, Rinzel J. 2011. Role of active dendritic conductances in subthreshold input integration. J Comput Neurosci 31:13–30. doi:10.1007/s10827-010-0295-7

Riedner BA, Vyazovskiy VV, Huber R, Massimini M, Esser S, Murphy M, Tononi G. 2007. Sleep Homeostasis and Cortical Synchronization: III. A High-Density EEG Study of Sleep Slow Waves in Humans. Sleep 30:1643–1657. doi:10.1093/sleep/30.12.1643

Schulz JM, Oswald MJ, Reynolds JNJ. 2011. Visual-Induced Excitation Leads to Firing Pauses in Striatal Cholinergic Interneurons. Journal of Neuroscience 31:11133–11143. doi:10.1523/JNEUROSCI.0661-11.2011

Schulz JM, Reynolds JNJ. 2013. Pause and rebound: sensory control of cholinergic signaling in the striatum. Trends in Neurosciences 36:41–50. doi:10.1016/j.tins.2012.09.006

Sela Y, Vyazovskiy VV, Cirelli C, Tononi G, Nir Y. 2016. Responses in Rat Core Auditory Cortex are Preserved during Sleep Spindle Oscillations. Sleep 39:1069–1082. doi:10.5665/sleep.5758

Sharott A, Doig NM, Mallet N, Magill PJ. 2012. Relationships between the Firing of Identified Striatal Interneurons and Spontaneous and Driven Cortical Activities In Vivo. Journal of Neuroscience 32:13221–13236. doi:10.1523/JNEUROSCI.2440-12.2012

Smith Y, Raju DV, Pare JF, Sidibe M. 2004. The thalamostriatal system: a highly specific network of the basal ganglia circuitry. Trends Neurosci 27:520–527. doi:10.1016/j.tins.2004.07.004 S0166-2236(04)00232-2 [pii]

Song WJ, Surmeier DJ. 1996. Voltage-dependent facilitation of calcium channels in rat neostriatal neurons. J Neurophysiol 76:2290–2306.

Stern EA, Jaeger D, Wilson CJ. 1998. Membrane potential synchrony of simultaneously recorded striatal spiny neurons in vivo. Nature 394:475–478. doi:10.1038/28848

Tanimura A, Du Y, Kondapalli J, Wokosin DL, Surmeier DJ. 2019. Cholinergic Interneurons Amplify Thalamostriatal Excitation of Striatal Indirect Pathway Neurons in Parkinson’s Disease Models. Neuron 101:444–458.e6. doi:10.1016/j.neuron.2018.12.004

Tanimura A, Lim SAO, Aceves Buendia J de J, Goldberg JA, Surmeier DJ. 2016. Cholinergic Interneurons Amplify Corticostriatal Synaptic Responses in the Q175 Model of Huntington’s Disease. Front Syst Neurosci 10. doi:10.3389/fnsys.2016.00102

Tchumatchenko T, Newman JP, Fong M, Potter SM. 2013. Delivery of continuously-varying stimuli using channelrhodopsin-2. Front Neural Circuits 7. doi:10.3389/fncir.2013.00184

Thomas TM, Smith Y, Levey AI, Hersch SM. 2000. Cortical Inputs to m2-Immunoreactive Striatal Interneurons in Rat and Monkey. Synapse 37:252–261.

Thorn CA, Graybiel AM. 2010. Pausing to Regroup: Thalamic Gating of Cortico-Basal Ganglia Networks. Neuron 67:175–178. doi:10.1016/j.neuron.2010.07.010

Threlfell S, Lalic T, Platt NJ, Jennings KA, Deisseroth K, Cragg SJ. 2012. Striatal Dopamine Release Is Triggered by Synchronized Activity in Cholinergic Interneurons. Neuron 75:58–64. doi:10.1016/j.neuron.2012.04.038

Tiroshi L, Goldberg JA. 2019. Population dynamics and entrainment of basal ganglia pacemakers are shaped by their dendritic arbors. PLoS Comput Biol 15:e1006782. doi:10.1371/journal.pcbi.1006782

Tubert C, Murer MG. 2021. What’s wrong with the striatal cholinergic interneurons in Parkinson’s disease? Focus on intrinsic excitability. Eur J Neurosci 53:2100–2116. doi:10.1111/ejn.14742

Ulrich D. 2002. Dendritic Resonance in Rat Neocortical Pyramidal Cells. Journal of Neurophysiology 87:2753–2759. doi:10.1152/jn.2002.87.6.2753

Wilson C, Chang H, Kitai S. 1990. Firing patterns and synaptic potentials of identified giant aspiny interneurons in the rat neostriatum. J Neurosci 10:508–519. doi:10.1523/JNEUROSCI.10-02-00508.1990

Wilson CJ. 2005. The mechanism of intrinsic amplification of hyperpolarizations and spontaneous bursting in striatal cholinergic interneurons. Neuron 45:575–585.

Wilson CJ. 2004. Basal Ganglia In: Shepherd GM, editor. The Synaptic Organization of the Brain. New York: Oxford University Press. pp. 361–413.

Wilson CJ, Goldberg JA. 2006. Origin of the slow afterhyperpolarization and slow rhythmic bursting in striatal cholinergic interneurons. J Neurophysiol 95:196–204. doi:00630.2005 [pii] 10.1152/jn.00630.2005

Yamanaka K, Hori Y, Minamimoto T, Yamada H, Matsumoto N, Enomoto K, Aosaki T, Graybiel AM, Kimura M. 2018. Roles of centromedian parafascicular nuclei of thalamus and cholinergic interneurons in the dorsal striatum in associative learning of environmental events. J Neural Transm 125:501–513. doi:10.1007/s00702-017-1713-z

Yarom O, Cohen D. 2011. Putative Cholinergic Interneurons in the Ventral and Dorsal Regions of the Striatum Have Distinct Roles in a Two Choice Alternative Association Task. Front Syst Neurosci 5. doi:10.3389/fnsys.2011.00036

