## Supplementary material for "Non-uniform distribution of dendritic nonlinearities differentially engages thalamostriatal and corticostriatal inputs onto cholinergic interneurons": Figure 3-figure supplement 1

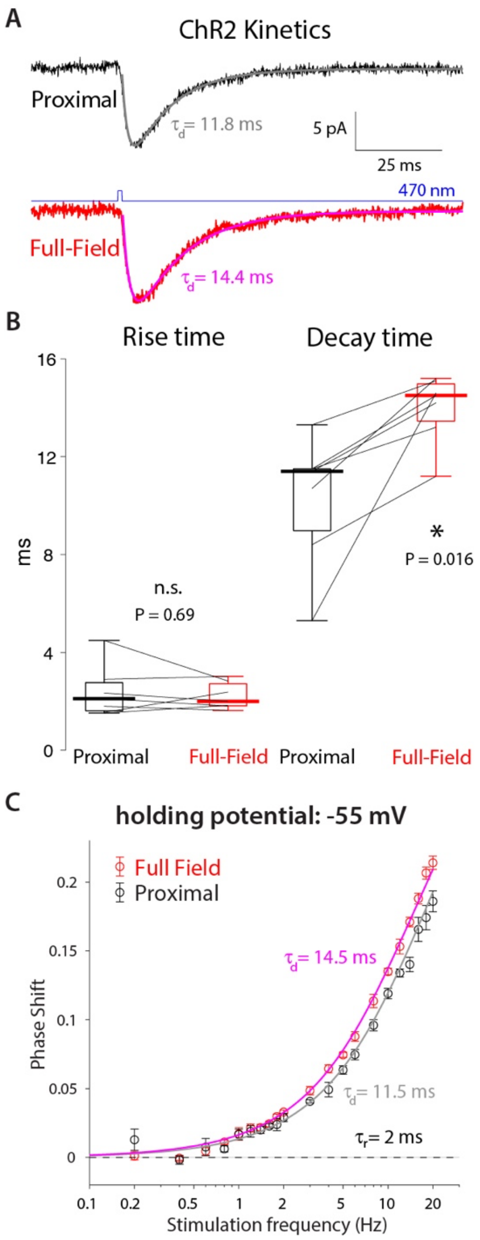


**Figure 3–figure supplement 1. High frequency phase delays in response to optogenetic activation are attributable to ChR2 kinetics and dendritic delays.** **A.** Mean somatic current responses to a 1 ms-long proximal (black, alpha-function fit in gray) and full-field (red, fit in magenta) 470 nm illumination. **B.** Distributions of rise and decay time constants (n = 7 CINs, N = 3 mice) demonstrate that the decay time constant was significantly larger by approximately 3 ms for the full-field illumination, due to dendritic delays. C. Phase delays of an alpha function with the observed time constants fit the empirical phased delays observed at -55 mV.
