## Supplementary material for "Non-uniform distribution of dendritic nonlinearities differentially engages thalamostriatal and corticostriatal inputs onto cholinergic interneurons": Figure 3-figure supplement 2

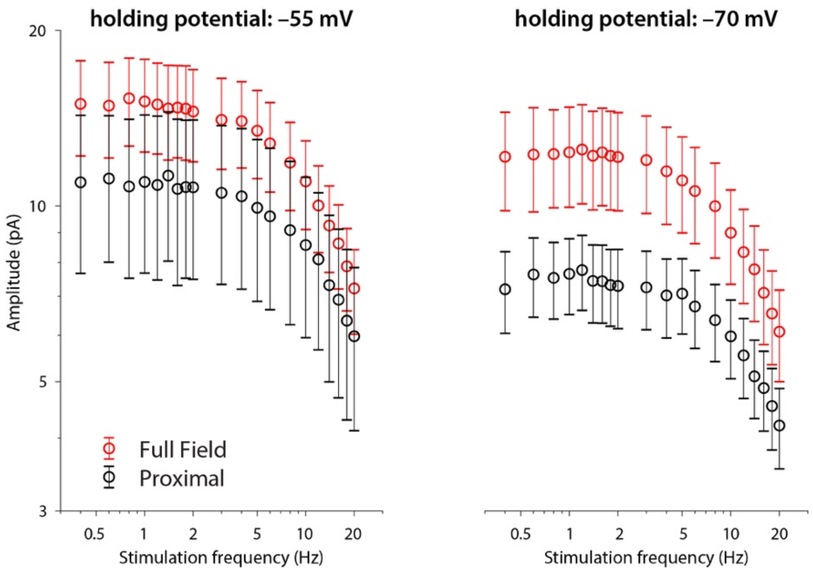


**Figure 3–figure supplement 2. Amplitude responses to proximal (black) and full-field (red) illumination at the two holding potentials.** While the curves at –70 mV hint at the presence of a low frequency resonance, the large error bars provide minimal constraints to model fits.
