## Supplementary material for "Non-uniform distribution of dendritic nonlinearities differentially engages thalamostriatal and corticostriatal inputs onto cholinergic interneurons": Figure 5-figure supplement 1

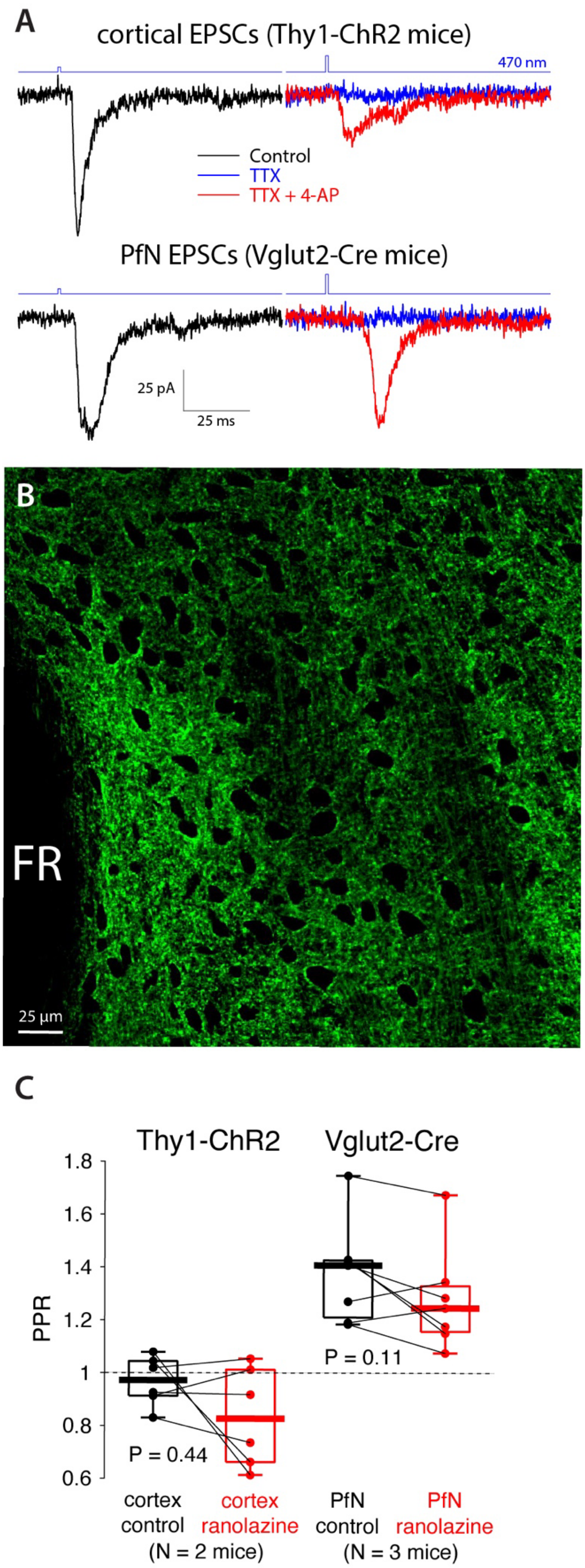


**Figure 5–figure supplement 1. Monosynaptic excitatory cortical and thalamic paired pulse ratios (PPRs) are not affected by ranolazine. A.** Optogenetic EPSCs in all CINs recorded in Thy1-ChR2 ($n = 8$ neurons, $N = 2$ mice) and in 6/7 CINs from Vglut2-cre mice, whose PfN was inoculated with AAVs harboring floxed ChR2 ($N = 2$) were monosynaptic, as 4-AP (100 µM) rescued release in the presence of TTX (1 µM). **B.** Coronal slice via the PfN in a Thy1-ChR2-EYFP mouse express EYFP in fibers only but not in somata, ruling out that intrastriatal optogenetic activation in these mice recruits thalamic inputs. FR – fasciculus retroflexus. **C.** Optical PPRs (100 ms interval) recorded in Thy1-ChR2 mice (left) or in Vglut2-cre mice whose PfN was transfected with AAVs harboring ChR2 (right) are unchanged by ranolazine.
