## Appendix 1 for "Non-uniform distribution of dendritic nonlinearities differentially engages thalamostriatal and corticostriatal inputs onto cholinergic interneurons"

### Appendix 1 – Fitting the quasi-linear model and opsin-mediated attenuation and phase shifts

<u>Modeling somatic voltage perturbations</u>

The properties of a quasi-linear membrane are captured by the following equations (Eqs. 4 in the Materials and Methods)

$$\alpha(f) = \gamma_R + \frac{\mu_n}{1+(2\pi f \tau_n)^2} + \frac{\mu_h}{1+(2\pi f \tau_h)^2}$$
$$\beta(f) = 2\pi f \left[\tau - \frac{\mu_n \tau_n}{1+(2\pi f \tau_n)^2} - \frac{\mu_h \tau_h}{1+(2\pi f \tau_h)^2}\right]$$

The membrane time constant is given by $\tau$ ; the total nonlinear conductance relative to leak is given by $\gamma_R$; the negative amplifying parameter is given by $\mu_n$, and the positive resonance parameter is given by $\mu_h$. $\tau_n$ and $\tau_h$ are the corresponding time constants of these nonlinear conductances. The notation and formalism used for describing the quasi-linear membrane is based on previous publications by us and others, where the values of the above parameters are extracted from the underlying biophysical models of the nonlinear conductances (Goldberg et al., 2007; Remme and Rinzel, 2011; Tiroshi and Goldberg, 2019). The reader is referred to those articles for further reading.

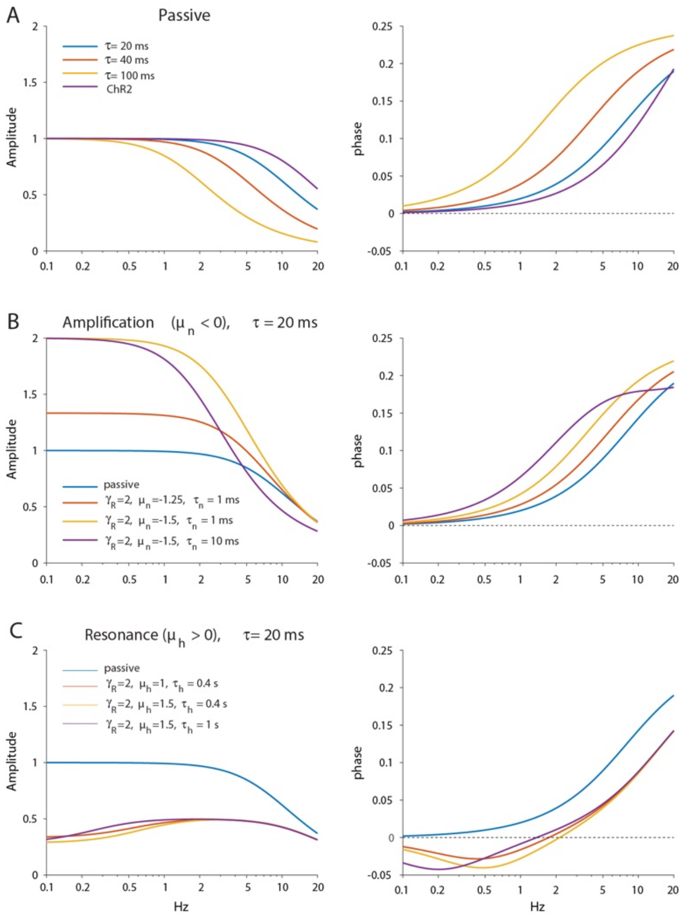


**Appendix 1–figure 1. Dependence of amplitude and phase responses on parameters of the quasi-linear model. A.** Passive dendrites are controlled by a single parameter $\tau$. Amplitude and phase responses for typical values of the empirical alpha function used to model the ChR2 response ($\tau_r = 2$ ms, $\tau_d = 11.5$ ms) are is shown in purple for comparison. **B.** Adding amplification increases the low frequency amplitude and phase response. **C.** Adding resonance reduces the low frequency amplitude response and introduces negative delays in the low frequency phase response.

We begin with the passive membrane, where $\mu_n = \mu_h = 0$ and $\gamma_R = 1$. In this case the amplitude, $(\alpha^2 + \beta^2)^{-1/2}$, and phase responses $\phi_s$ (of Eq. 5 in Materials and Methods), are those of a low-pass (LP) filter controlled by a single parameter $\tau$. As $\tau$ is increased the cut-off frequency of the LP filter becomes smaller and the phase shift increases in the lower frequencies (Appendix 1–figure 1A).

Next we introduce the amplifying current. We immediately see the boosting of the lower frequencies in the amplitude response as $\mu_n$ is made more negative. Here too, increasing $\tau_n$ shifts the cut-off to lower frequencies and increases the phase shift in the lower frequencies (Appendix 1–figure 1B). Note that the amplifying current can only increase the phase shifts.

When introducing a restorative current a resonance is created. While the resonance can be observed in the amplitude response as the formation of a non-zero maximal amplitude, *it is much more robustly* observed in the negative lobe that forms in the phase response at low frequencies (Appendix 1–figure 1C). Increasing $\mu_h$ deepens the negative lobe and shifts the zero-crossing (and the resonance peak) to a higher frequency. In contrast, increasing $\tau_h$ shifts the zero crossing (and resonance) to a lower frequency. Note that for self-consistency when adding a nonlinear conductance, $\gamma_R$ must also be increased above 1 (Goldberg et al., 2007; Remme and Rinzel, 2011).

Modeling dendritic optogenetic perturbation

When modeling the optogenetic activation of the dendrite we form a cascade of two filters: that of the channelrhodopsin-2 (ChR2, Eq. 6 in Materials and Methods, Appendix Figure 1A, "purple") and that of the dendrite. The parameters of ChR2 kinetics are based on our measurement of the rise ($\tau_r$) and decay ($\tau_d$) time constants (of the alpha function) that we fit to the data (Figure 3–figure supplement 1). However, the fact that the estimate of $\tau_d$ is systematically larger for the full-field illumination relative for the proximal illumination, must result from the cable properties of the dendritic arbor (Tiroshi and Goldberg, 2019). This means that the underlying decay kinetics must be shorter than the decay time recorded under both proximal and full-field illumination. Therefore, for fitting the dendritic model, we chose $\tau_d = 10$ ms and $\tau_r = 0.2$ ms as representative values, both of which are in agreement with the literature on ChR2 kinetics in expression systems (Nagel et al., 2003).

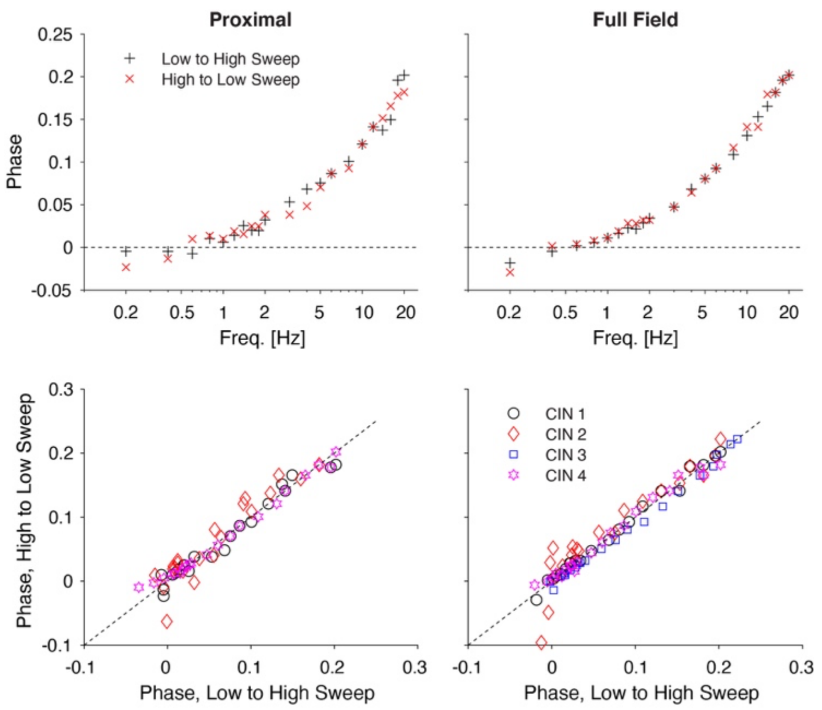


**Appendix 1–figure 2. Reversing the optogenetic frequency sweep does not affect the phase estimates**. Top: estimates of the phases as a function of frequency for one neuron for increasing (black) and decreasing (red) frequencies for proximal and full-field illumination. Bottom: Scatter plot of phases recorded for decreasing vs. increasing frequencies (for 4 cells), show that the values cluster around the diagonal. Holding potential -70 mV.

An additional concern about the ChR2 kinetics is that they may have a longer time scale effect on our estimates of the amplitude and phase responses. To alleviate this concern we compared the estimates attained when sweeping the frequencies from low-to-high to sweeping them from high to low, and found that they do not differ (Appendix 1–figure 2).

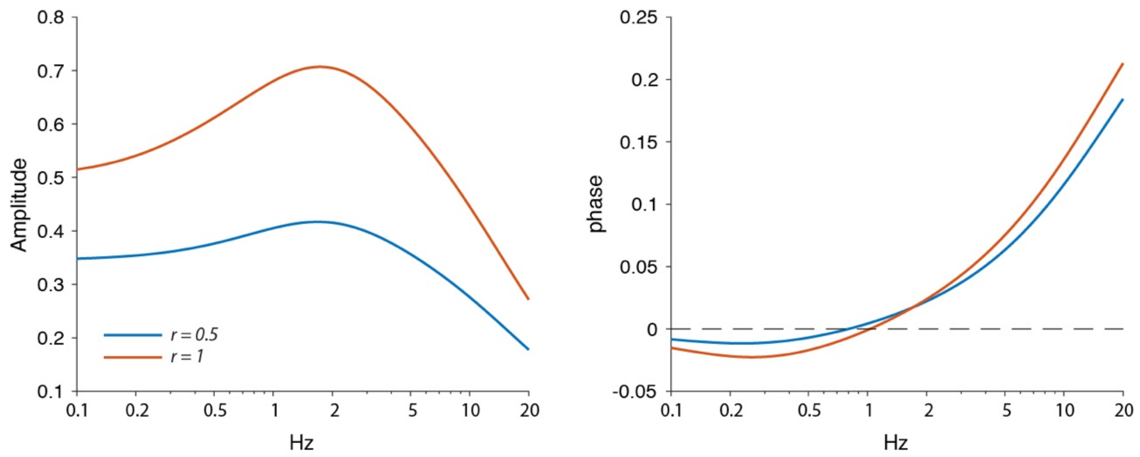


**Appendix 1–figure 3. Amplitude and phase response arising from the ChR2 kinetics and a quasi-linear dendrite.** For a homogeneous distribution of quasi-linear properties results in a stronger resonance when a larger portion of the dendrite (*r=1 vs. r=0.5*) is illuminated, which can be seen as a sharper peak and a negative phase region with more negative phases. Other parameters: $\gamma_R = 2$, $\tau = 20$ ms, $\mu_n = -1.5$, $\mu_h = 2$, $\tau_n = 1$ ms, $\tau_h = 0.8$ s, $\tau_r = 0.2$ ms, $\tau_d = 10$ ms.

The dendritic amplitude response is given by $(1 - e^{-pr})(p^2 + q^2)^{-1/2}$ (Tiroshi and Goldberg, 2019) where *r* is the length of the dendrite being illuminated (in units of dendritic space constants, Appendix 1–figure 3A). Illumination of a longer extent of the dendrite results in a larger phase shift in the higher frequencies of the phase response (Appendix 1–

figure 3B), as seen in the present study (Figure 3C,D), and in our previous study of dendrites of GABAergic neurons in the substantia nigra pars reticulata (Tiroshi and Goldberg, 2019). Importantly, for a given positive value of $\mu_h$, when more of the dendrite is illuminated the negative lobe becomes larger (Appendix 1–figure 3B), in contrast to what we found in the case of the cholinergic interneurons (CINs, Figure 3D). This mathematical fact, strengthens our conclusion that in order to recapitulate the empirical observation of a smaller negative lobe (Figure 3D), the density of the restorative current (e.g., the HCN current) must decrease when more of the dendritic membrane is being illuminated.

Our model thus contains 6 free parameters ( $\tau_r$ and $\tau_d$ were not treated as free parameters, as explained above) for the somatic perturbations (Figures 1&2) and one additional parameter $r$ for the optogenetic dendritic perturbations (Figure 3) (see Table 1 in main text). Of course, the whole purpose of the model is to provide a more generalized method to characterize the impact of nonlinearities on dendritic integration, thereby reducing the number of free parameter to only this handful. Despite the significant dimensional reduction achieved by neglecting the fine details of the biophysics and kinetics of each of the channels that give rise to the nonlinearity (as well as the detailed dendritic morphology), we nonetheless attain several free parameters, which could support multiple viable fits to the same data, and might lead to overfitting of the data. To address this issue, we thus approached the model fitting as follows.

First, because a) the phase response can be fit more robustly than the amplitude response (the phase response does not depend strongly on the intensity of the stimulus); b) the negative phase region is a more robust measures of resonances than the amplitude (e.g, compare the left and right panels in Figure 1C); and c) the error bars on the phase are much tighter - we fit our parameters based solely on the phase responses (In Figures 1C & 2B,D, Figure 3–figure supplement 2). Consequently, we fit a single parameter for the amplitude of the impedance curves using the parameters of the fit attained from the phase shifts.

Second, we only included parameters when they were necessary. Based on the empirical data demonstrating that (TTX-sensitive) NaP currents produce no negative lobes in the phase response, but nevertheless boost responses at -55 mV, we fit those data (e.g., Figure 2B) with a single amplifying nonlinearity. Only for the HCN currents that empirically gave rise to the negative lobe (e.g., Figures 2D & 3D) did we use the full model (i.e., 6 and 7 free parameters, respectively).

Third, we conducted least-square curve-fitting (Matlab, Mathworks) by restricting the values of the parameters so that the reflect physiologically reasonable values, particularly for the membrane, NaP and HCN time constants (Goldberg et al., 2007) as follows:

| Parameter | $\mu_n$ | $\mu_h$ | $\tau_n$ (ms) | $\tau_h$ (s) | $\gamma_R$ | $\tau$ (ms) | $r$ |
|---|---|---|---|---|---|---|---|
| min | -5 | 0 | 0.1 | 0.01 | 1 | 10 | 0 |
| max | 0 | 5 | 100 | 10 | 10 | 100 | 2 |
| initial guess | -1 | 1 | 20 | 1 | 2 | 50 | 1 |

The two following qualitative findings gave us confidence in our approach, and in our conclusions regarding the dendritic distribution of nonlinearities. First, the *r* extracted from the full field configuration was always larger than the *r* extracted for the proximal illumination, indicating that the measurements were sensitive to whether only a proximal region of the CIN was illuminated or its entire dendritic field. Second, the $\mu$'s extracted for the full field illumination were smaller in absolute magnitude than those extracted for the proximal illumination, strengthening the conclusion that the channels that give rise to them are expressed primarily proximally.

In summary, while the quasi-linear model gives rise to multiple parameters, it is still a significantly dimensionally-reduced representation of dendritic morphology and filtering. Careful use of parameter fitting can be used to extract the qualitative properties of the dendrites, including regarding the localization of nonlinearities.
