## Appendix 2 for "Non-uniform distribution of dendritic nonlinearities differentially engages thalamostriatal and corticostriatal inputs onto cholinergic interneurons"

### Appendix 2 – Boosting of proximal *vs.* distal inputs in a model of a nonlinear dendrite

The dendrite was modeled (in XPPAUT) by discretizing the following nonlinear cable equation (with $dx = 0.1$ and $dt = 0.25$ ms).

$$\tau \frac{\partial}{\partial t} V(x,t) = \lambda^2 \frac{\partial^2}{\partial x^2} V(x,t) - [V(x,t) - V_L] - \gamma_n n(x,t)[V(x,t) - V_{Na}] H(\lambda - x) + \tau A \delta(t - t_0)\delta(x - x_0)$$

$$\tau_n \frac{\partial}{\partial t} n(x,t) = n^{\infty}[V(x,t)] - n(x,t)$$

where $0 < x < 5\lambda$ with boundary conditions $\frac{\partial}{\partial x} V(0,t) = \frac{\partial}{\partial x} V(5\lambda,t) = 0$ and $V_L = -60$ mV, $V_{Na} = 50$ mV, $\tau = 50$ ms, $\tau_n = 1$ ms, $\gamma_n = 0.6$ and $n^{\infty}(V) = \left[1 + e^{-(V+45)/3}\right]^{-1}$. $H(x)$ is the Heaviside function, indicating that only the region where $0 < x < \lambda$ expresses the NaP current. Proximal stimulation is at $x_0 = \lambda$ and distal stimulation is at $x_0 = 2\lambda$, and $t_0 = 25$ ms. To simulate a passive dendrite, we set $\gamma_n = 0$. The resting membrane potential –60 mV ($=V_L$) in the passive dendrite and –59.7 mV in the nonlinear dendritic region.

The figure below (Appendix 2–figure 1) depicts the voltage at the soma ($x = 0$) in response to proximal (solid line) and distal (dashed line) stimulation pulses at three intensities, where the minimal intensity (Appendix 2–figure 1A) was consequently multiplied by 2 and then by 4 (Appendix 2–figure 1B&C, respectively). Responses in the passive dendrite are depicted in black, whereas the responses in the nonlinear dendrite are indicated in red. In Appendix 2–figure 1D, we plotted the ratio of the amplitude in response to the stimulation, normalized to the response to the weakest stimulus in the passive dendrite (black lines in panel Appendix 2–figure 1A). The black symbols indicate that in the case of the passive dendrite, the amplitude of the response scales exactly linearly with the amplitude of the stimulation pulse for both proximal and distal stimulation.

The red symbols indicate the case of the nonlinear cable. For the weakest stimulus, the model replicates the quasi-linear approximation, and exhibits a slight amplification relative to the passive cable (i.e., the normalized amplitude is slightly larger than 1), and this amplification is *identical for proximal and distal inputs* (i.e., the plus sign and circle overlap), even though the NaP is restricted to the proximal dendrite. Indeed, this will hold true for the quasi-linear approximation, as well, because it is a linear model. However, when stronger stimuli are used, it becomes evident that the proximal stimulation is amplified (nonlinearly) more than the distal input. The reason for this is that the distal perturbation decays considerably before it arrives at the boosting region. Therefore, it is much less boosted than the proximal perturbation that engages the nonlinearity of the NaP current.

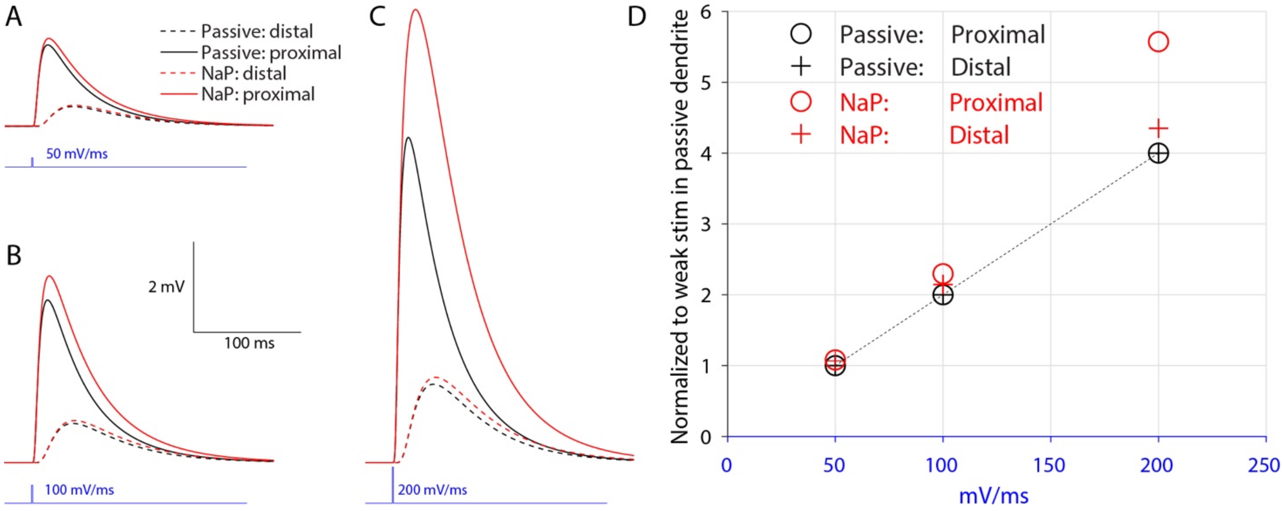


**Appendix 2–figure 1: Non-linear boosting of proximal and distal inputs in a cable model where the amplifying nonlinearity is restricted to the proximal region**. **A**. Somatic ($x = 0$) voltage perturbations elicited by stimulating either a proximal region (solid line) that expresses the nonlinearity or a distal region (dashed line) that does not. Red traces are in the full-nonlinear cable and the black traces are in the case of a passive (linear) dendrite. **B**. same as A only with stimulations that are twice as large. **C**. same as A only with stimulations that are four times as large. **D**. Amplitude of responses normalized to the response in the weakest condition in a the passive dendrite (i.e., solid black lines in panel A). Because the passive dendrite is linear all responses scale perfectly linearly (dotted black line), and proximal and distal inputs decay precisely to the same degree (the black crosses overlap the black circles precisely). In the case of the nonlinear dendrite, in the weakest stimulation (panel A) some boosting can be observed (red marks are slightly above 1). However, both distal and proximal inputs are boosted to the same degree, indicating that the system behaves as expected from a quasi-linear amplifying membrane). For stronger stimulations, the nonlinearity amplifies the proximal input (where the nonlinearity is expressed) more than it amplifies the distal input.
